# NovaClone: A Network-Based Algorithm for Clonal and Subclonal Genotyping of Barcoded Transgene Integrations

**DOI:** 10.64898/2026.05.11.724244

**Authors:** Sebastian Prillo, Dana Rimini, Pedro Olivares-Chauvet, Yun S. Song, Nir Yosef

## Abstract

Single-cell lineage tracing technologies are providing new and powerful ways to interrogate the evolution and divergence of cell populations in cancer, development, and other contexts. A key initial step in any such analysis is the grouping of cells into clonal populations, based on clone-level marks. Unfortunately, clone calling is prone to technical effects due to sequencing errors, missing data, multiplets, background noise, and accidental sharing of clonal barcodes between unrelated clones (homoplasy). We present `NovaClone`, a principled algorithm for hierarchical clone calling that is broadly applicable to all current tracing technologies, including both static barcoding and the more recent evolving tracers. We benchmark `NovaClone` on simulated and real data to show that it outperforms the current solutions in terms of both quality and speed, thereby helping to mitigate one of the most prevalent problems with single-cell lineage tracing. To complement `NovaClone`, we introduce a suite of algorithm-agnostic quality control metrics to evaluate clone calls when ground truth is not available. `NovaClone` and the associated QCs are available through the open source Python package `nova-clone`.

## 1 Introduction

Single-cell lineage tracing technologies have emerged as powerful tools to study growing cell populations *in vivo* [1–7]. At a basic level, these technologies work by randomly integrating into the DNA of an initial pool of progenitor cells a set of lineage tracing reporter genes, using common recombinant DNA techniques such as Piggybac transposition or lentiviral infection. Each integrated gene is designed to include a unique region that serves as a barcode (referred to as an *integration barcode*). Each progenitor cell in the pool thus holds a random set of integration barcodes, which will be inherited by all its descendants. Profiling the barcode sequences of each descendant cell is commonly done with single cell RNA-sequencing (as the reporter regions are transcribed), but it can also be done through DNA sequencing or fluorescent-based methods (for non-random barcodes) [8]. Once available, these barcodes can help reveal the clone of origin of each cell and thus enable clone-wise analysis downstream.

Lineage tracing approaches differ by the extent of subclonal associations that can be read off the data. Some studies rely on static barcoding through a single round of integration of reporter genes that stay fixed, providing information only about founding lineages and not about internal sub-clonal structure [1–7]. Other studies utilize several successive rounds of integration to reveal splits into subclones that took place during these rounds [9]. While these integration events provide discrete checkpoints, emerging studies also incorporate dynamic recording, where lineage tracing reporter genes are randomly and continuously mutated throughout the experiment, usually through designated CRISPR/Cas9 target sites [1–3, 5–7, 9], enabling the analysis of fine subclonal relationships between the captured cells. The need for clone calling, however, is shared by all types of studies.

A simple example of the barcode integration process is illustrated in Figure 1. In this example there are two rounds of integrations: In the first round, one progenitor cell from our pool acquires the integration barcodes (or `intBCs` for short) {*AAA, BBB*} and another acquires the barcodes {*EEE, FFF*} . (Note: for simplicity of exposition, we use the entire English alphabet to describe these DNA barcodes). Cells are then allowed to recover before a second round of integration. During this period, cells may continue to divide. A second round of integration is performed, where each subclone inherits its parental barcodes and acquires additional ones, leading to subclonal populations carrying barcodes such as {*AAA, BBB, CCC*},{*AAA, BBB, DDD*},{*EEE, FFF, GGG*}, and {*EEE, FFF, DDD, HHH*} . Note that not all subclones may acquire new integrations during the second round. Also note how the barcode *DDD* is acquired by two distinct subclones - this is referred to as *homoplasy* and is one of the many factors that will complicate the task of clone calling. Once the cells have been engineered with these barcodes, they can be transferred to *in vivo* models or *in-vitro* cultures where they continue to grow.

**Fig. 1.**
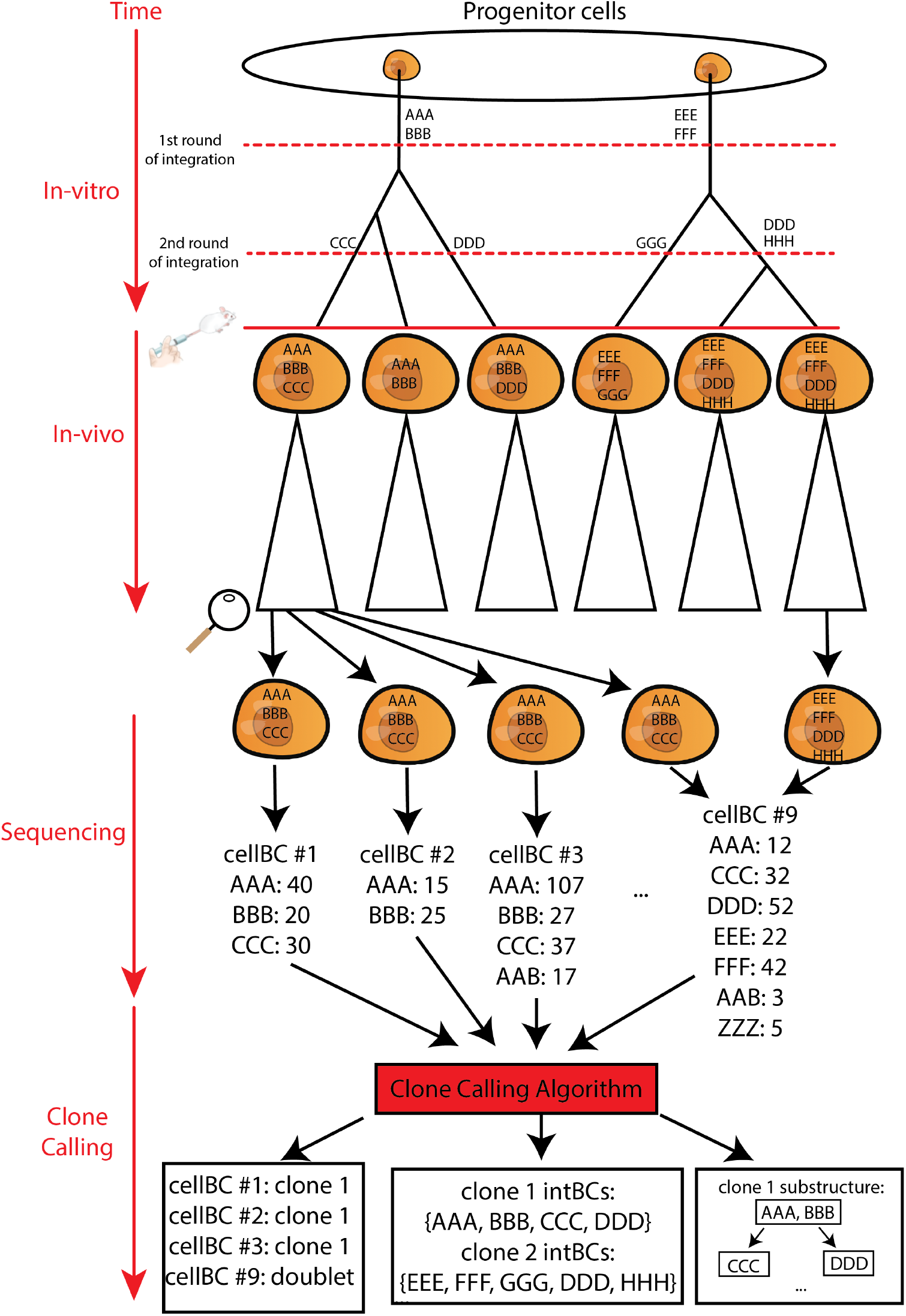
Complete lineage tracing pipeline. Progenitor cells undergo two rounds of integration during which cell division occurs, creating a subclonal structure. The cells are then sequenced, producing an integration barcode table that may be impacted by inaccuracies such as missing data, sequencing errors and multiplets. The clone calling algorithm then assigns cells to clones (identifying multiplets in the process), identifies the barcode set of each clone, and reconstructs subclonal structures.

By the end of the experiment, the cells are harvested and profiled (e.g., with single-cell RNA-sequencing), where the sequence for each `intBC` and the number of molecules of its respective mRNA (referred to here as Unique Molecular Identifier count, or `UMI_count` for short) can be evaluated for each cell (identified uniquely by a barcode sequence, denoted as `cellBC`). The task of clone calling is to infer the grouping of cells into clones from tables that consist of these three features (`cellBC`, `intBC`, `UMI_count`), where each row corresponds to a distinct (`cellBC`, `intBC`)-pair. These tables, however, often include inaccuracies that arise through the process of reading out the integration barcodes, whether using sequencing of cells in droplets, or through fluorescent imaging. Typical issues include missing data, background RNA, or (for the former type of approaches) sequencing errors and empty droplets. Multiplets are also an important caveat, with two or more cells captured by the same cell barcode (in sequencing-based approaches) or due to segmentation issues (in spatial, imaging-based technologies).

The current approaches for calling clones are usually based on simple clustering of cells based on similarities of their integration barcodes [4, 7]. This approach, however, may suffer from *over-merging*, whereby two unrelated sets of cells are reported to be clonally related; this may occur due to homoplasy, undetected multiplets, background RNA, and sequencing errors misleading the algorithm. It may also suffer from *under-merging*, whereby two sets of clonally related cells are reported to originate from different clones, e.g., due to sequencing errors or missing data. These errors can have a large impact on downstream analysis, e.g., in estimating the numbers and sizes of clones, their rates of growth, within-clone heterogeneity of gene expression, and other clone-level properties. At present, however, relatively little attention has been devoted to the clone calling problem, and the details about how this procedure is carried out (e.g., parametrization) and the related quality control (QC) metrics are often lacking in published studies.

In this work, we introduce `NovaClone`, a principled algorithm for the clone calling problem which is broadly applicable across lineage tracing technologies, as well as a set of QC metrics to assess the quality of any clone call. The only major assumption of our algorithm is that the number of shared `intBCs` (i.e. homoplasy degree) between two distinct clonal populations is at most one. Sharing of more than one `intBC` is extremely unlikely by design for most lineage tracing technologies. For example, if the `intBCs` are randomized DNA sequences of length 14, then there are 4^14^ possible barcodes. The probability that two clones share more than one `intBC` by chance is small, even if the probability of synthesis across barcode sequences is non-uniform. `NovaClone` is a network-based method, and leverages concepts that have been previously successful in analyzing sequencing data, such as the UMITools algorithm [10]. In addition, we provide a panel of generally applicable QC metrics and plots to certify the high quality of the resulting clone calls, or otherwise report potential issues. Our panel of QC metrics is *agnostic* to the details of the clone calling algorithm, as it uses only the raw data and the inferred clone call. It is thus applicable to the output of any clone calling algorithm, not just ours. In fact, we use our QC metrics to highlight issues with the clonal quality in previously reported works.

Taken together, our broadly applicable `NovaClone` clone calling algorithm and set of QC metrics set a new standard for clone calling, which promises to improve the quality and rigor of all single-cell lineage tracing experiments.

## 2 Methods

The input to `NovaClone` is a table consisting of triplets of (`cellBC`, `intBC`, `UMI_count`) listing the number of unique occurrences of an `intBC` in a given `cellBC`, as is typical of single-cell readouts. The output is a map assigning each `cellBC` to its respective clone (or set of clones, in the case of multiplets), as well as the coarse tree describing the subclonal structure of each clone (Figure 1). At a high level, `NovaClone` consists of two main phases (Figure 2):

1. **Phase 1-Constructing clonal signatures**: A unique set of `intBCs` characterizing each clone – which we call *clonal signatures* – are constructed. These signatures have the property that they do not overlap between clones and include only high-confidence `intBCs` (using criteria that we describe next). This is the most involved phase of the algorithm.
2. **Phase 2-Mapping** `cellBCs` **to clones and calling full** `intBC` **sets**: In this phase, `cellBCs` are mapped to their clones, based on overlaps with the clonal signatures. Once all `cellBCs` have been assigned, the complete set of `intBC` in each clone is evaluated. This step helps recover lower abundance `intBCs` which may not have made it into the initial clonal signatures.

**Fig. 2.**
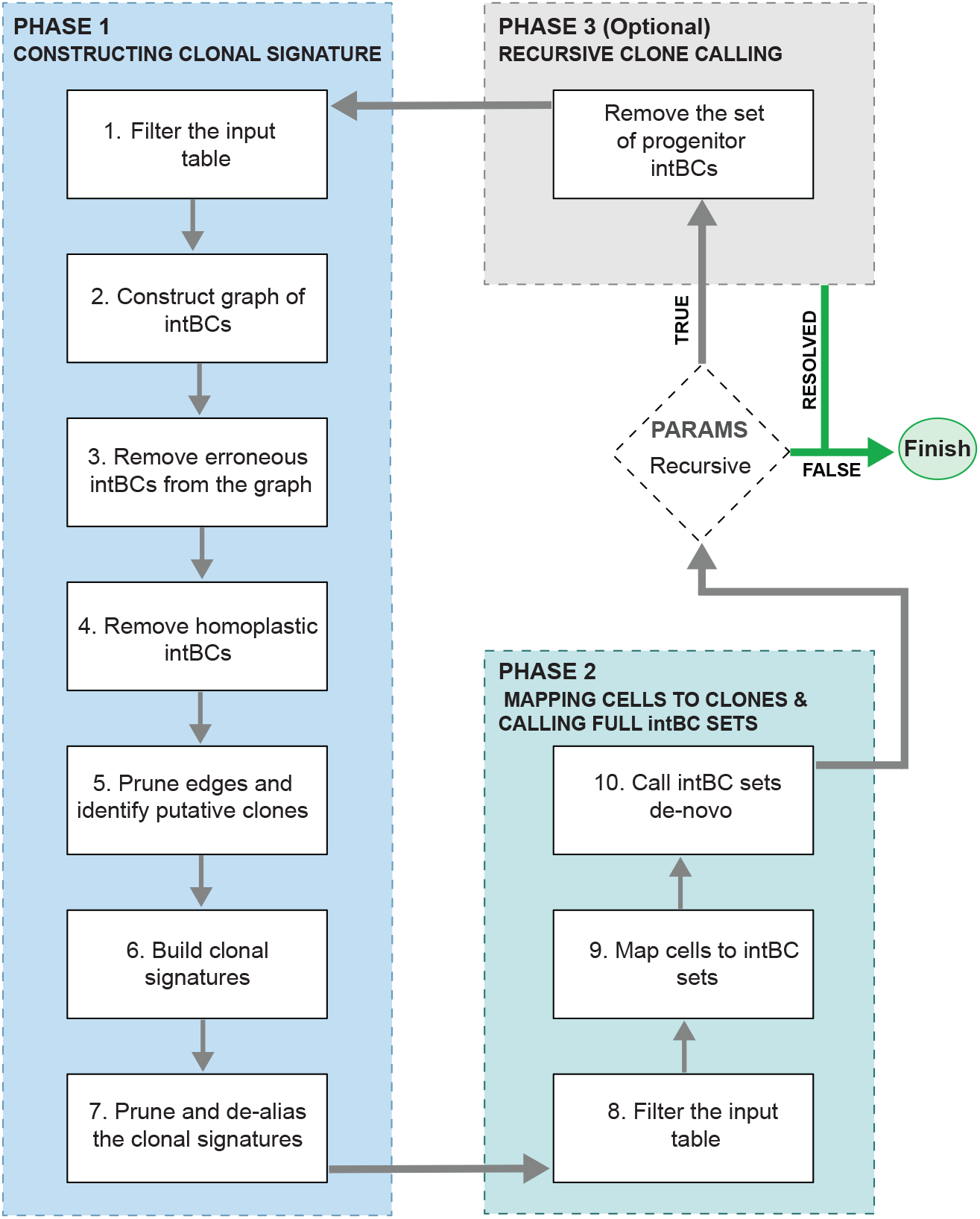
Flow chart of NovaClone. The `NovaClone` algorithm consists of two main phases: phase 1 constructs *clonal signatures* and phase 2 maps `cellBCs` to clones using these clonal signatures and finally calls `intBC` sets de novo. When run with the *recursive* option, `NovaClone` runs recursively to identify subclonal structures.

**Fig. 3.**
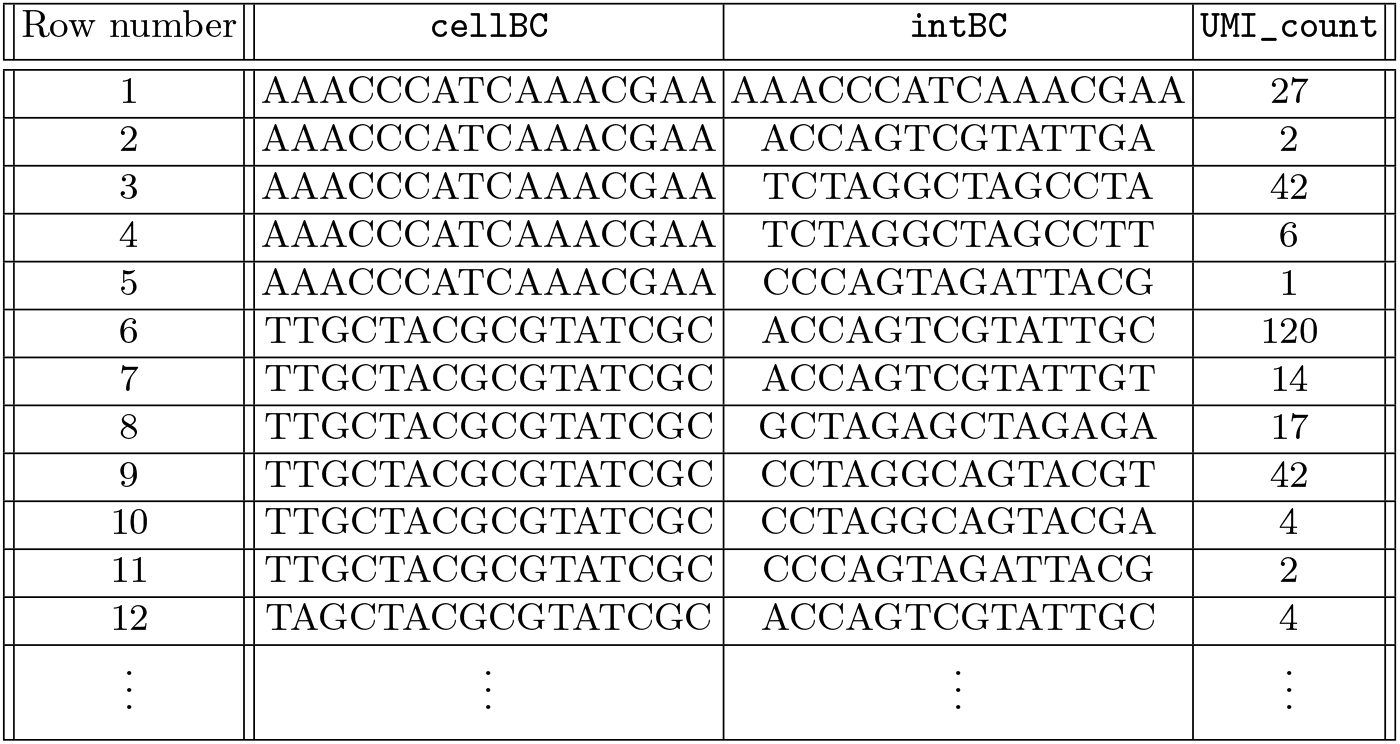
Example table including rows with common issues in lineage tracing technologies that `NovaClone` addresses, including sequencing errors in `intBCs` (rows 2, 4, 7, 10) and `cellBCs` (row 12), as well as background RNA contamination, where an `intBC` from another cell (rows where the `intBC` is CCCAGTAGATTACG) is incorrectly associated with additional `cellBC`.

Formally, the input to `NovaClone` is a table with columns: (`cellBC`, `intBC`, `UMI_count`). We call this the *intBC table*. A realistic example of an *intBC table* is as follows:

### 2.1 Phase 1: Construct clonal signatures

The goal of the first phase of `NovaClone` is to enumerate the clones in the dataset and determine a *clonal signature* for each clone, namely - a set of `intBCs` that uniquely identify it. To this end, we require that clonal signatures be pairwise disjoint between clones. Thereafter, a `cellBC` will be associated with a clone if and only if it contains at least one `intBC` in its respective signature. Clonal signatures should therefore include only non-homoplastic integrations. In the toy example of Figure 1, valid clonal signatures would be {*AAA, BBB*} and {*EEE, FFF*}, which mark the two major clones. Additionally, subclonal integrations that are non-homoplastic may also serve to expand the clonal signatures and reduce the number of non-assigned cells. In the case of our toy example, valid expanded clonal signatures would be {*AAA, BBB, CCC*} and {*EEE, FFF, GGG, HHH*} respectively. The barcode *DDD* should not be part of any clonal signature as it is homoplastic. Similarly, sequencing errors and background RNA should also be excluded from clonal signatures. The construction of clonal signatures is conducted in several steps, described below.

#### Filtering spurious entries from the input table

Our first step is to identify and discard any entries (rows) in the input table that are likely to be noisy, either in the identity of the `cellBC`, the `intBC`, or the association between them. We first remove entries that are supported by a small number of UMIs (`UMI_count`). Such a count filter is a standard practice in scRNA-seq [11] and helps discard sequencing errors (both on `intBCs` and `cellBCs`) and background RNA, which tend to appear in small quantities. By default, our algorithm will discard all rows in the input tables where the `UMI_count` is less than 10, but this parameter can be tuned according to the numbers of reads in the sample of interest. For example, when we run `NovaClone` on the *macsGESTALT* dataset [7], we used a lower threshold of two UMIs due to the overall lower molecular counts per cell, compared to later assays. More generally, we recommend to set the UMI cutoff by considering a histogram of UMI counts across all rows in the input table and use the leftmost mode (in case the count distribution is multi-modal) or a low quantile (otherwise).

Next, we remove all rows corresponding to `intBCs` which are observed in less than a certain number of `cellBCs`. We use a threshold of 10 as default. This accomplishes the goal of removing rare `intBCs` which are thus less helpful for identifying clones. The cutoff of 10 provides certain appealing theoretical guarantees, which we discuss later for identifying both (i) sequencing errors and (ii) spurious edges in the (to be defined) `intBC` graph. Finally, our implementation of `NovaClone` allows the user to provide a *custom filtering function* with which to filter rows from the input table. Since in most assays `intBCs` lengths are fixed by design, this filter can be used to remove erroneous `intBC`. For instance, in the example in Figure 1 all `intBCs` should be 14bp in length and therefore the entry in row 1 will be removed. Depending on how reads were processed, custom filters can be designed to capture other suspicious entries, e.g., ones that include `intBC` sequences with unresolved bases, similarity to primer sequences, or low quality scores. While the user could filter these manually before calling `NovaClone`, we strove to make the algorithm self-contained. While errors can still remain after these first filtering steps, the count filters applied provide the statistical power needed for the sequencing error and homoplasy detection steps that follow.

#### Constructing a graph of integration barcodes

Following the work of UMITools [10], a graph representation of `intBCs` (nodes) and the similarities between them (edges), provides a convenient tool for denoising the data, and ultimately inferring clonal signatures. In this graph, an edge connects two nodes if there exists at least one `cellBC` that is associated with both. Each node *x* is assigned with a weight *c*(*x*), set to the number of `cellBCs` in which it appears. Each edge (*x, y*) is assigned with a weight *c*(*x, y*), set to the number of `cellBCs` in which both `intBCs` occur. An important quantity is the *co-occurrence ratio* of two `intBCs` *x* and *y*, which we define as:

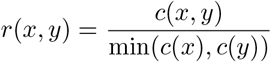

Equivalently, letting:

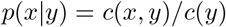

be the probability of observing `intBC` *x* when `intBC` *y* is present in the `cellBC`, then we have *r*(*x, y*) = max(*p*(*x* | *y*), *p*(*y* | *x*)). Intuitively, *r*(*x, y*) is ‘the maximum probability with which one of the two barcodes implies the other’.

Figure 4 illustrates what the `intBC` graph might look like for the example in Figure 1.

**Fig. 4.**
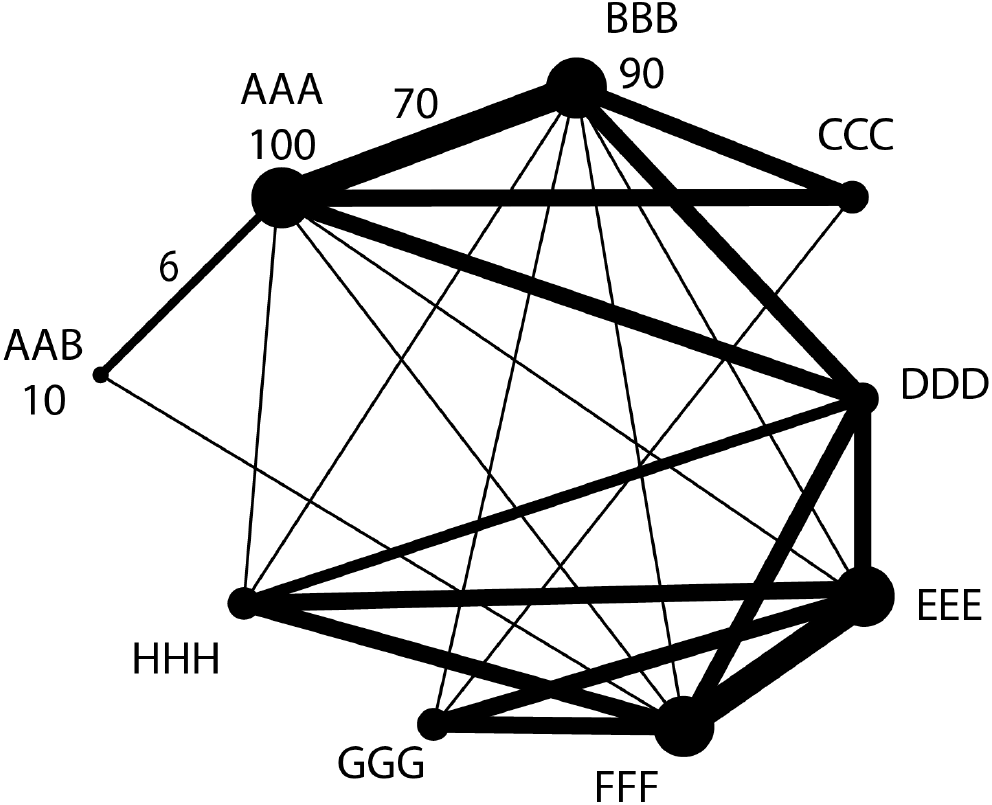
The intBC graph. In the `intBC` graph, there is one node per `intBC`. The *weight* or *counts c*(*x*) of node *x* is the number of `cellBCs` which the `intBC` appears in. For each pair of `intBCs` which co-occur in some `cellBC`, we add an edge with *weight* or *counts c*(*x, y*) equal to the number of `cellBCs` in which both `intBCs` co-occur. In this example, we have that *c*(*AAA*) = 100, *c*(*BBB*) = 90, *c*(*AAB*) = 10, *c*(*AAA, AAB*) = 6, *c*(*AAA, BBB*) = 70. The *ratio* is a key quantity which specifies the maximum probability with which one barcode implies the other, for instance 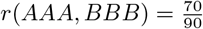.

#### Removing erroneous integration barcodes from the graph

While a UMI filter can be effective in mitigating the effect of sequencing errors, some may still persist. This can happen, for instance, in the case of well-captured `intBCs` (e.g. row six in our example) where even a small fraction of wrongly processed reads can be sufficient to pass the UMI filter (as it is defined on the absolute number of molecules observed). The `intBC` graph provides an intuitive way to further identify and discard these `intBCs`.

The key observation is that `intBCs` which are sequencing errors tend to co-occur (in the same `cellBC`) with their correct version, with the latter observed at a higher count. Thus, we search the `intBCs` graph for `intBC` *x* which satisfy the following property: there exists another `intBC` *y* such that (1) *y* has more counts than *x*, (2) the edit distance between *x* and *y* is small (by default, ≤ 4), and (3) *p*(*y* | *x*) ≥ 0.5. These criteria support the notion that *x* and *y* originated from the same locus and that *y* is the correct version of the `intBC` sequence. We thus declare *x* to be a putative sequencing error and prune this node from the `intBC` graph. Notably, it is possible that `intBCs` which are not sequencing errors are pruned in this step. The goal of this step, however, is to achieve high recall for identifying sequencing errors, as false negatives lead to more significant errors downstream (e.g. overmerging of clones).

The default value of 0.5 should work well because at this point in the algorithm, each `intBC` is supported by at least 10 `cellBCs` (by virtue of the initial `min_intbc_cellbc_count` filter). Therefore, for a sequencing error to make it through, the worst-case scenario is that the sequencing error appears in just 10 `cellBC` and its correct version is only detected 4 of those 10 times, which is extremely rare. Indeed, let *p* be the probability that when capturing a sequencing error, the original sequence is also captured. Assuming *p* is equivalent to the sequencing saturation, which is typically 90% for high-quality libraries [12], we fail to filter the sequencing error if Bin(10, 0.90) ≤ 4. This event occurs with a probability of approximately 1.4 × 10^−4^ (roughly 1 in 7,000).

As above, the most effective edit distance cutoff may change between samples or lineage tracing technologies and it can therefore be tuned by users of `NovaClone`. A useful way to set this cutoff is to consider the distribution of edit distances between every pair of `intBCs` in the same `cellBC` prior to any filtering. We expect this distribution to be multi-modal and to show a left peak due to sequencing errors. Figure S10 shows an example of this on real data, demonstrating a clear bimodality, with a left peak at 2-3 bp. Figure 5 shows the `intBC` graph after pruning sequencing errors. Since *AAB* and *AAA* frequently co-occur (with a ratio of *r*(*AAA, AAB*) = 0.6) and their edit distance is small, then *AAB* is flagged as a potential sequencing error and removed from the `intBC` graph.

**Fig. 5.**
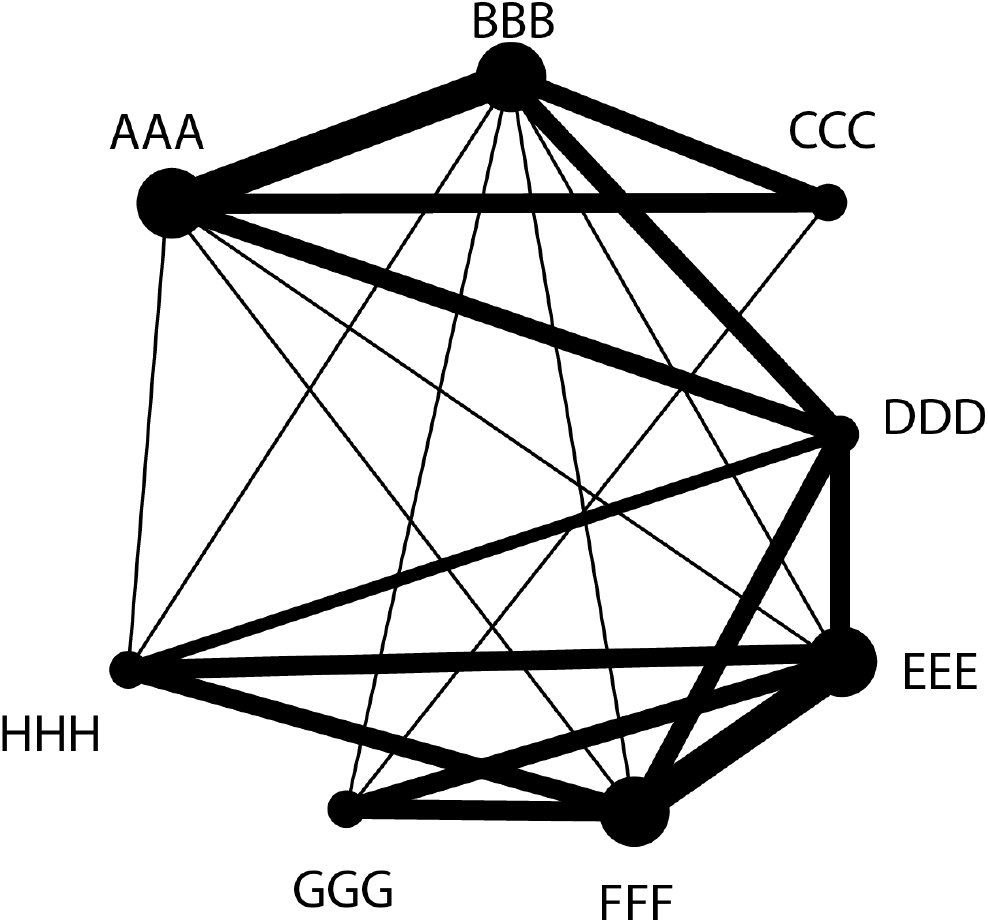
The intBC graph after pruning sequencing errors. By using statistical filters, `NovaClone` prunes sequencing errors with high sensitivity. This `intBC` graph of our running example is shown after pruning of sequencing errors.

#### Removing putative homoplastic integration barcodes caused by synthesis bias and background RNA contamination

An `intBC` is *homoplastic* if it was incorporated in two or more independent integration events (i.e., it is present in more than one clone or subclone). In Figure 1, `intBC` *DDD* is homoplastic. While longer `intBC` sequences make it less likely for homoplastic `intBCs` to exist, the process of `intBC` DNA synthesis is not entirely random [13–16] and so some barcode sequences are more likely to be synthesized than others. Alternatively, background RNA contamination may cause a single `intBC` to be captured across many cells in the assay. To devise a criterion for identifying these putative homoplastic cases, first consider the ideal case of no missing data, no multiplets, no sequencing errors, and at least two `intBC` integrations per progenitor cell. In that case, all `cellBCs` in which a given non homoplastic `intBC` occurs will also include all other `intBCs` in the progenitor cell of the clone (irrespective of whether those other `intBCs` are homoplastic); for example, in Figure 1, the non-homoplastic `intBC` *CCC* always appears together with *AAA* (and *BBB*). Thus, if we define the *co-occurrence weight* for each `intBC` *x* as

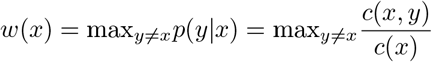

then the co-occurrence weight will be exactly 1 for non-homoplastic `intBCs` and likely lower than 1 for homoplastic `intBCs` as homoplastic `intBCs` appear with different sets of `intBCs`. For example, in Figure 1, the homoplastic `intBC` *DDD* sometimes co-occurs with *AAA, BBB, CCC* (clone 1) and sometimes co-occurs with *EEE, FFF, GGG, HHH* (clone 2). Therefore, *w*(*x*) *<* 1 would accurately predict homoplastic `intBCs` in an ideal, noiseless setting.

Due to technical artifacts such as missing data, the weight *w*(*x*) can frequently drop below 1 for non-homoplastic `intBCs`. For example, in Figure 4, neither *AAA* nor *BBB* are homoplastic, yet their co-occurrence weights are *w*(*AAA*) = 0.7 and *w*(*BBB*) = 70*/*90. To account for this noise, we use a relaxed threshold of *w*(*x*) *<* 0.5.

We do not expect this relaxed criterion to detect all homoplastic `intBCs`: for instance, an `intBC` appearing in a large and a small clone will not be detected. However, this step is effective at detecting and removing `intBCs` which appear in many clones, e.g. due to synthesis bias or background RNA contamination, which makes our algorithm robust to such effects. In our running example, `intBC` *DDD* is homoplastic as it emerges in two unrelated subclones. Whether or not it will be pruned at this step has to do with the numbers of cells that were captured from the relevant clones. We will assume that *DDD* is **not** pruned in this example; later steps of the algorithm will identify *DDD* as homoplastic. However, for our algorithm to succeed, it is crucial that *after* this pruning step no two clones share more than two `intBC` sequences, such as in our running example (Figure 1).

At this point of the algorithm, the filtered `intBC` graph provides a conservative view, which focuses on nodes (`intBCs`) that are less likely to reflect sequencing errors (i.e., they are sufficiently distinguishable and are supported by a minimal number of observations). Furthermore, with the last homoplasy pruning step, we further assume that the chances are low for any remaining `intBC` to appear in more than two clones. For those remaining `intBCs` that do appear in two clones, we assume that the chances are low for any pair of them to appear in the exact same two clones, since this would involve coordinated occurrence of four relatively rare events. Therefore, **we assume that from now on, no two clones share more than one** `intBC`. If this assumption is violated, our algorithm will incorrectly merge the involved clones, resulting in an *over-merging* error.

#### Pruning spurious edges

Having aggressively pruned sequencing errors from our `intBC` graph, we are confident that remaining `intBCs` represent true integrations. However, co-occurrence of two `intBCs` (i.e. *c*(*x, y*) *>* 0) does not imply a shared clonal origin. This is due to multiplets, whereby a single `cellBC` captures `intBCs` from multiple cells leading to spurious co-occurrences of `intBCs`. Given that multiplets are a common issue in both droplet- and imaging-based sequencing technologies, this stage in the algorithm is designed to address them.

Formally, call an edge (*x, y*) in the `intBC` graph *faithful* if *x* and *y* co-occur in the same clone. Otherwise, call the edge (*x, y*) *spurious*. In our running example, edges such as (*AAA, BBB*), (*AAA, CCC*), (*CCC, DDD*), (*DDD, GGG*) are all faithful, whereas (*AAA, EEE*) is spurious. Our goal in this step is to prune from the `intBC` graph all edges that are spurious, with a high level of sensitivity. To identify spurious edges with high sensitivity, note that if (*x, y*) is spurious, then very likely *c*(*x*), *c*(*y*) *>> c*(*x, y*) (i.e. *r*(*x, y*) is close to zero). Indeed, assuming without loss of generality that *c*(*x*) *> c*(*y*), then for every cell in the original sample in which `intBC` *y* is expressed and captured: (i) if this cell is captured by a unique `cellBC`, it will contribute one count to *c*(*y*) and none to *c*(*x, y*), (ii) *only* if this cell is part of a multiplet **and** that multiplet is against a cell expressing `intBC` *x*, will we observe a count for *c*(*y*) and *c*(*x, y*). Since case (i) is statistically much more likely than (ii) (which involves two low-probability events), we will thus likely have *r*(*x, y*) ≈ 0. In contrast, for faithful edges (*x, y*), whenever *y* is observed, we will likely observe *x* (unless it has been epigenetically silenced). Thus, a threshold of *r*(*x, y*) *<* 0.5 should have high sensitivity for pruning spurious edges.

The only issue from a statistical point of view is that there are many pairs (*x, y*) to test, so if we are not careful, some spurious edges might pass through our filter (false negatives). This is the point of the algorithm in which it is crucial that we have only retained `intBCs` with high counts (*c*(*x*) ≥ 10 by default). This ensures that for each pair (*x, y*) that is spurious, the probability of passing the cutoff is vanishingly small. For example, if there are 10 clones of the same size and the multiplets rate is 20%, then for fixed `intBCs` *x* and *y* in different clones: for every `cellBC` expressing *y* that we observe, we only expect 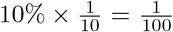 of the time for *y* to co-occur with *x*. Thus, the probability of spurious edge (*x, y*) passing our cutoff is 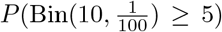 which is less than 1 in a million, so even with just *c*(*x*) ≥ 10 we obtain vanishingly small probability of failure. In practice most `intBCs` have more than 10 counts, so these probabilities are even smaller. Taken together, even with multiple testing considered, this step of our algorithm prunes *all* spurious edges with high probability, leaving only faithful ones.

In our running example, Figure 6 shows the result of applying this pruning step to the `intBC` graph: (i) all spurious edges such as (*AAA, EEE*), (*CCC, HHH*) are removed, (ii) faithful edges corresponding to parent-descendant integrations such as (*AAA, BBB*), (*AAA, CCC*), (*DDD, AAA*), (*DDD, DDD*) will be retained, (iii) faithful edges corresponding to non-parent-descendant integrations such as (*CCC, DDD*), (*GGG, DDD*), (*GGG, HHH*) will also be removed. Overall, the graph is highly sparsified. We are almost ready to extract clonal signatures.

**Fig. 6.**
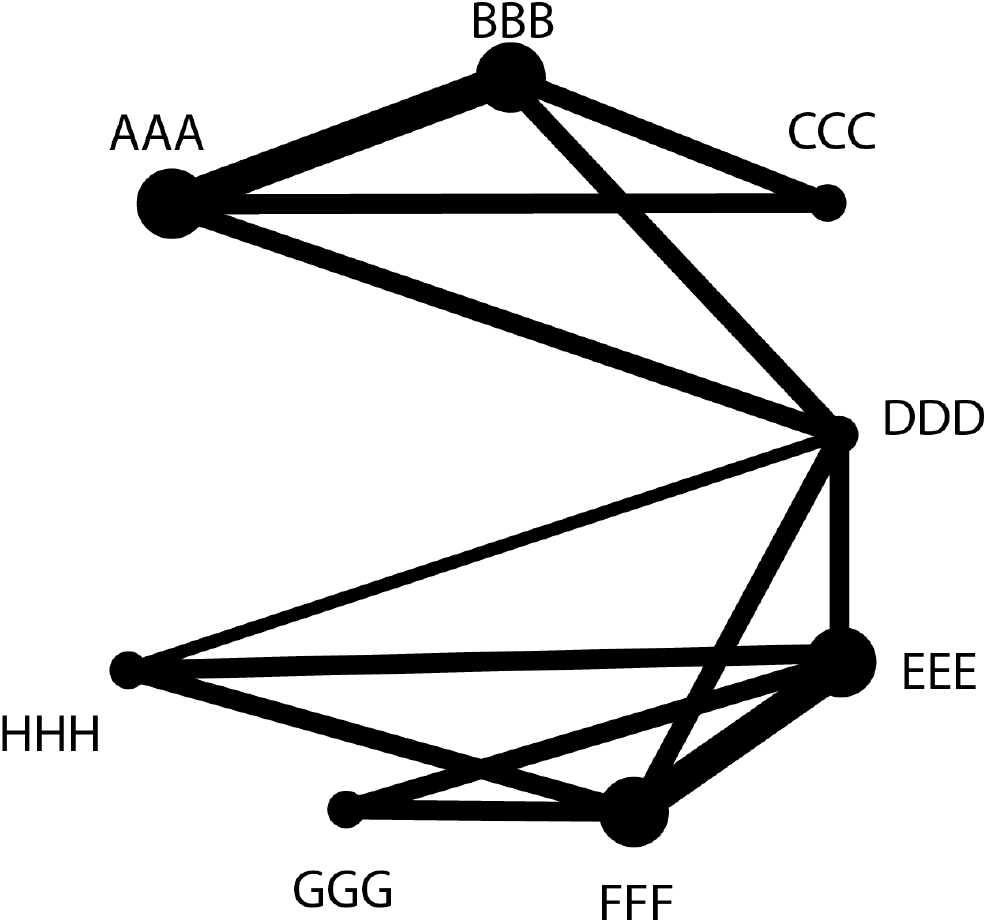
The intBC graph after the last pruning step. The `intBC` graph in our running example after pruning spurious edges. The graph is significantly more sparse and its clonal structure starts to become apparent. Note how the homoplastic intBC *DDD* still remains, forming a “bridge” between the two major clones .

#### Building preliminary clonal signatures

At this point of the algorithm, all edges (*x, y*) in the `intBC` graph correspond to `intBCs` *x* and *y* that co-occur in some same clone. Moreover, since we assume that at this point no two clones may share more than one `intBC` through homoplasy, it follows that *x* and *y* co-occur in *only one clone*. Thus, as our next step, we partition the pruned graph into clusters, where different clusters may share at most one `intBC`. With our assumptions, discarding such “bridge” nodes (representing `intBCs` that are shared by two clones) will result in a partition of the graph into disjoint sets, each representing some clone (see Figure 6). In our running example, node *DDD* is such a “bridge” (Figure 6).

To achieve this decomposition into clusters which share at most one `intBC`, we first identify all maximal cliques in the `intBC` graph. Call these sets of `intBCs` *K*_1_, *K*_2_, …, *K*_*p*_. Next, if two sets *K*_*i*_ and *K*_*j*_ intersect in at least two elements, we replace them by *K*_*i*_ ∪ *K*_*j*_. We perform this operation until there are no two sets of `intBCs` that overlap in more than one `intBCs`. Let the final collection of `intBC` sets be *K*_1_, *K*_2_, …, *K*_*p*_*′* . We consider this as our preliminary enumeration of clones in the data and refer to the set of `intBCs` in each candidate clone as its putative clonal signature. In Figure 7 we show the result of running this algorithm on our ongoing example.

**Fig. 7.**
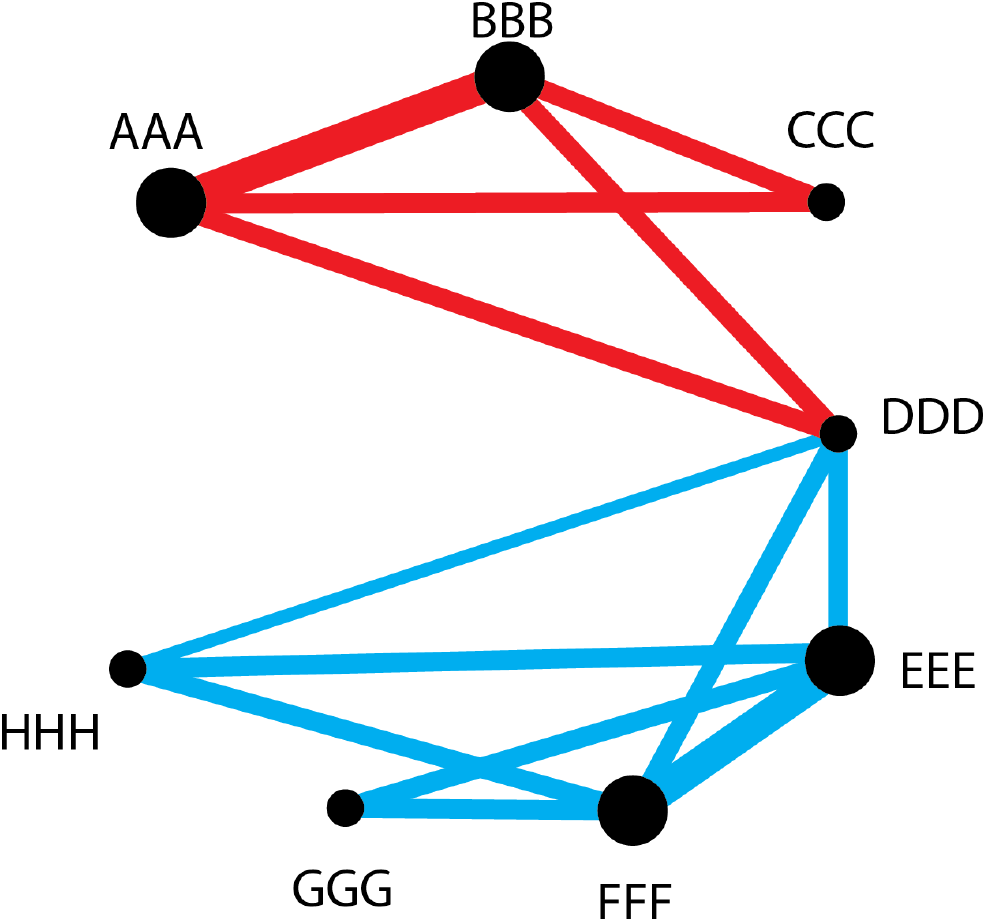
The coresets of the intBC graph identify clones. The partitioning of the `intBC` graph into putative clones starts with detection of maximal cliques. In this case {*AAA, BBB, CCC*}, {*AAA, BBB, DDD*}, {*DDD, EEE, FFF, HHH*}, and {*DDD, EEE, FFF, GGG*} . The clonal signatures are repeatedly checked for overlap and merged if their overlap is at least 2 `intBCs`. In our example, the cliques {*AAA, BBB, CCC*} and {*AAA, BBB, DDD*} get merged into {*AAA, BBB, CCC, DDD*} and the cliques {*DDD, EEE, FFF, HHH*} and {*EEE, FFF, GGG*} get merged into {*DDD, EEE, FFF, GGG, HHH*} . At this point, no more merges are possible, so these are determined to be our preliminary clonal signatures. Note that the preliminary clonal signatures {*AAA, BBB, CCC, DDD*} and {*DDD, EEE, FFF, GGG, HHH*} intersect on *DDD*, which is not enough to merge. It is in this step that the crucial assumption that **two clones should not share more than one** `intBC` must hold, otherwise those two clones will risk being merged together, as their preliminary signatures will be merged.

There are two final issues to take care of: (1) it is possible that two of the `intBC` sets *K*_*i*_ and *K*_*j*_ still represent the same clone, and (2) there may be homoplastic `intBCs` in these sets. The main reason for (1) is that pruning spurious edges might also prune some faithful edges due to high but imperfect specificity (e.g. (*CCC, DDD*) gets pruned in our example). This leads to subclonal integrations forming smaller disjoint cliques that still correspond to the same progenitor clone. We deal with these two issues in the next steps.

#### Pruning and de-aliasing the preliminary clonal signatures

A remaining concern which we address next is the possible existence of pairs of `intBC` sets (*K*_*i*_, *K*_*j*_) which represent the same clone. To tackle this, we consider the association between `cellBCs` and putative clones (i.e., putative clonal signatures) and look for pairs of putative clones that are associated with a similar set of `cellBCs`. Specifically, we map a `cellBC` to a putative clone if it intersects its putative clonal signature in at least 2 `intBCs`. We then search for pairs *K*_*i*_ and *K*_*j*_ for which (1) *K*_*i*_ is associated with fewer `cellBCs` than *K*_*j*_, (2) more than 50% of the `cellBCs` associated with *K*_*i*_ also map to *K*_*j*_. In that case, we remove *K*_*i*_ from the set of preliminary clonal signatures. In our running example, no aliased signatures were found.

As a final step of this first phase of `NovaClone`, we further clean the clonal signatures in two ways: (1) We enforce a lower bound on the size of clones by discarding signatures with fewer than 30 associated cells. We do this because very small clones are usually not of interest in studies. However, this threshold can be changed if one desires to identify smaller clones. Note that since we have already enforced *c*(*x*) ≥ 10, all clonal signatures will have at least 10 cells to support it. (2) We make the clonal signatures disjoint by removing any `intBC` that still participate in more than one signature. In our running example, `intBC` *DDD* is removed from the putative clonal signatures, yielding {*AAA, BBB, CCC*} and {*EEE, FFF, GGG, HHH*} as the final clonal signatures. Note that these signatures correspond to the true `intBC` sets of each clone in Figure 1 after removing homoplastic intBCs, which is the optimal outcome.

### 2.2 Phase 2: Mapping cells to clonal signatures and calling full intBC sets

The outcome of the first phase of the algorithm was pairwise disjoint clonal signatures *K*_1_, *K*_2_, …, *K*_*q*_ that uniquely identify each clone. In this second phase, we use the clonal signatures to map `cellBCs` to clones (using a *de novo* approach) and then, based on the mapped `cellBCs`, derive the full `intBC` set for each clone de novo.

To this end, we return to the raw data as our input, namely the `intBC` table filtered by a requirement for minimal `UMI_count` (as in phase one). We map each `cellBC` to a clone if it intersects the clonal signature by at least two `intBCs`. A `cellBC` mapped to more than one clonal signature is called a multiplet, and `cellBCs` mapped to a unique clone are deemed unique single-cells. We then take each clone in turn, and consider the set of `intBCs` observed in at least one of its associated cells. This list can contain: (1) true `intBCs`, some of which can be homoplastic, (2) sequencing errors, (3) other `intBCs` with undesirable properties (e.g., related to sequence content, originally filtered with the custom filtering function). To clean this list, we traverse it from the `intBCs` that has the highest frequency (number of `cellBCs` it is observed in) to the lowest. We add an `intBC` to the clone’s full `intBC` set if: (1) there is no other higher frequency `intBC` at an edit distance lower than 4, (2) the `intBC` appears in at least a critical number of cells, taking as our cutoff by default: max(20%of cells in clone, 10) (3) it passes any custom filtering if provided. Notably, the 20% threshold is intended to retain high-quality, informative `intBCs` that appear in many cells within a clone. However, this threshold can be relaxed to retain `intBCs` present in only a small subset of cells, which is a characteristic of sub-clonal integrations much like *GGG* in Fig. 1.

### 2.3 Phase 3 (optional): Resolving subclonal structures via recursive clone calling

In some experimental designs, `intBCs` can be integrated into the progenitor population in multiple rounds, creating a rich subclonal structure. To resolve this structure, `NovaClone` allows for a recursive mode, in which clone detection is re-applied separately on each of the clones that were detected in the previous round. The main challenge is that naively applying `NovaClone` to cells from a single clone will always identify just one clone, since all these cells share the same progenitor `intBCs` from their common ancestor. To overcome this limitation, we use a three step approach: first, we *infer* the progenitor `intBCs` that define the parent clone (details below); second, we *remove* these shared `intBCs` from the data; and *finally*, we run `NovaClone` on the remaining `intBCs` to detect subclonal structure.

#### Remove the set of progenitor intBCs

To infer the set of progenitor `intBCs` of a clone, we start from its set of `intBCs` derived from the de-aliasing step. This is the clonal signature of the clone, plus homoplastic `intBCs`. By construction, the `intBC` graph induced by these `intBCs` is connected. We re-build the `intBC` graph for these `intBCs`, and proceed to pop nodes from the graph from highest count to lowest count until the graph disconnects. Once the graph disconnects, if {*x*_1_, *x*_2_, …, *x*_*k*_} are the `intBCs` we popped, then we declare them to be the `intBCs` of the progenitor cell.

In our running example, this means that for clone 1, we take the set `intBC` graph formed by {*AAA, BBB, CCC, DDD*} . We first pop *AAA* - the highest frequency `intBC`-but the graph does not disconnect (as *BBB* connects to *CCC* and *DDD*.) Therefore, we proceed to pop the next highest frequency `intBC`, in this case *BBB*. The `intBC` finally disconnects, revealing {*AAA, BBB*} as the progenitor `intBC` set - which agrees with the ground truth shown in Figure 1. For the second clone, *EEE* and *FFF* get popped and then the `intBC` graph disconnects.

Having identified the putative set of progenitor `intBCs` in that clone, we take the cells from that clone together with its inferred `intBC` set and *remove* the progenitor `intBCs`. In our running example, this means that for clone 1 we remove {*AAA, BBB*} from the clone’s `intBC` set, thus retaining only {*CCC, DDD*}. For clone 2, we retain only {*DDD, HHH, GGG*}.

#### Run NovaClone recursively

We proceed to call `NovaClone` recursively on this new data. In our example, the recursive call identifies two (sub)clonal signatures for clone 1: {*CCC*} and {*DDD*} . For clone 2, the two (sub)clonal signatures found are {*DDD, HHH*} and {*GGG*} . Since the algorithm is defined recursively, it can proceed to deeper levels of recursion (sub-sub clones etc.), per the user’s input.

Notably, (sub)clone calling is is a-priori easier at this stage, since the sets of `intBC` were filtered (for sequencing errors and any other custom filtering rules), and more importantly - since homoplasy *within* a single clone is rarer than in the entire dataset (simply due to the lower number of events). Therefore, when running `NovaClone` recursively we use more permissive parameters-allowing for clonal signatures of size 1, and requiring only one `intBC` of overlap for the merging procedure. Since the subclones are naturally smaller in size, we also allow a lower bound on the minimum number of cells required for calling sub-clones (10 by default). Finally, in cases where sub-clonal partition was detected, it might be the case that not all cells were associated to a sub-clone (since we require support by at least two `intBC`). To address this, we only accept the sub-clonal partitions if at least 70% of the cells have been associated.

## 3 Quality Control Metrics

After `NovaClone` runs (or *any* clone calling algorithm), one can perform checks to validate that the clone call is of high quality, or otherwise report abnormalities to the user. These QC metrics are algorithm-agnostic, and may therefore be used regardless of the clone calling method. All we require is that the clone call report the grouping of `cellBCs` into clones, the `intBC` set of each clone, and the final retained `intBCs` of each `cellBC`. For avoidance of confusion, in the case of `NovaClone`, these `intBC` sets are the final ones computed by the algorithm in phase 2 (as opposed to the clonal signatures computed in phase 1).

### 3.1 Multiplet rate

We define the **multiplet rate** *r*_multiplet_ as the fraction of `cellBC` that represent more than one cell. This rate is computed from the clone call by computing the fraction of `cellBC` mapped to more than one clone. This metric is particularly valuable because for some instruments the multiplet rate can be roughly estimated a-priori. For example, for a Chromium Next GEM Single Cell 3’ lane with 10000 cells loaded the expected multiplet rate is approximately 8% [17].

### 3.2 Percent unmapped cells

Another simple metric is the percent of `cellBCs` that were not mapped to any clone (*r*_unmapped_). We expect some `cellBCs` to not map to any clone because they correspond to empty droplets or clones that are too small to detect. If *r*_unmapped_ is large, this may indicate potential issues with the quality of the raw data or that of the data pre-processing routines (from *fastq* files to a barcode table).

### 3.3 Clone sizes and the numbers of integration barcodes

Focusing on the successfully mapped cells (the complementary set to the above two), additional QC metrics look into how large the clones are and how many `intBCs` are there per clone. The QC component of `NovaClone` plots these statistics for inspection by users. This allows the user a bird’s-eye view of the data, enabling the identification and removal of outliers - i.e., clones that passed our filters but are still much smaller than others or include very few `intBCs` per cell (Figure 8).

**Fig. 8.**
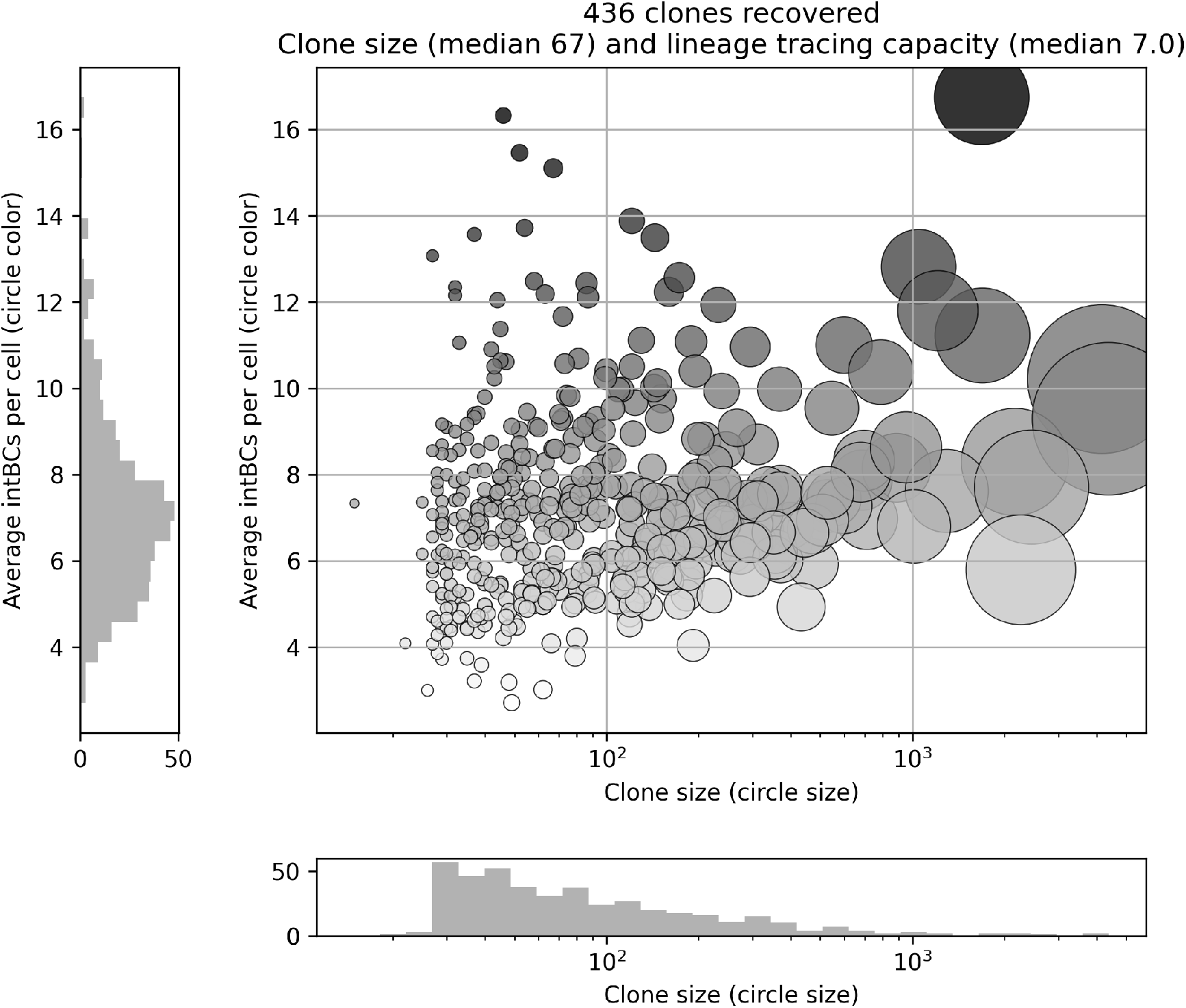
NovaClone Quality Control pipeline output: Clone sizes and lineage tracing capacities. Example taken from real data. Each point represents a clone. The x-axis shows clone size, also reflected by point size, and the y-axis shows the average number of `intBCs` per `cellBC` in the clone, encoded also by point color.

### 3.4 Intra-clonal kinship

Even randomly grouping the cells into clonal populations may produce a reasonable plot like the one above. Thus, we wish to evaluate whether the cells in each clonal population are indeed clonally related.

To this end, we define the *kinship* between two cells as the number of `intBCs` that they share, where we only focus on `intBCs` that are part of clone’s final `intBC` set (thereby ignoring sequencing errors, background RNA, and any other technical artifacts). The *intra-clonal kinship* is then defined as the distribution of kinships over all pairs. Notably, since in some studies the clones can be very large, if a clone has more than 200 cells, we randomly subsample 200 cells to compute its kinship distribution. The kinship distribution of each clone can be plotted as a row in a heatmap, as seen in Figure 9. The user can then choose to exclude clonal populations with a low intra-clonal kinship, as suspected erroneous clonal populations. By default, `NovaClone` warns the user about those clones with a mean intra-clonal kinship less than 2.

**Fig. 9.**
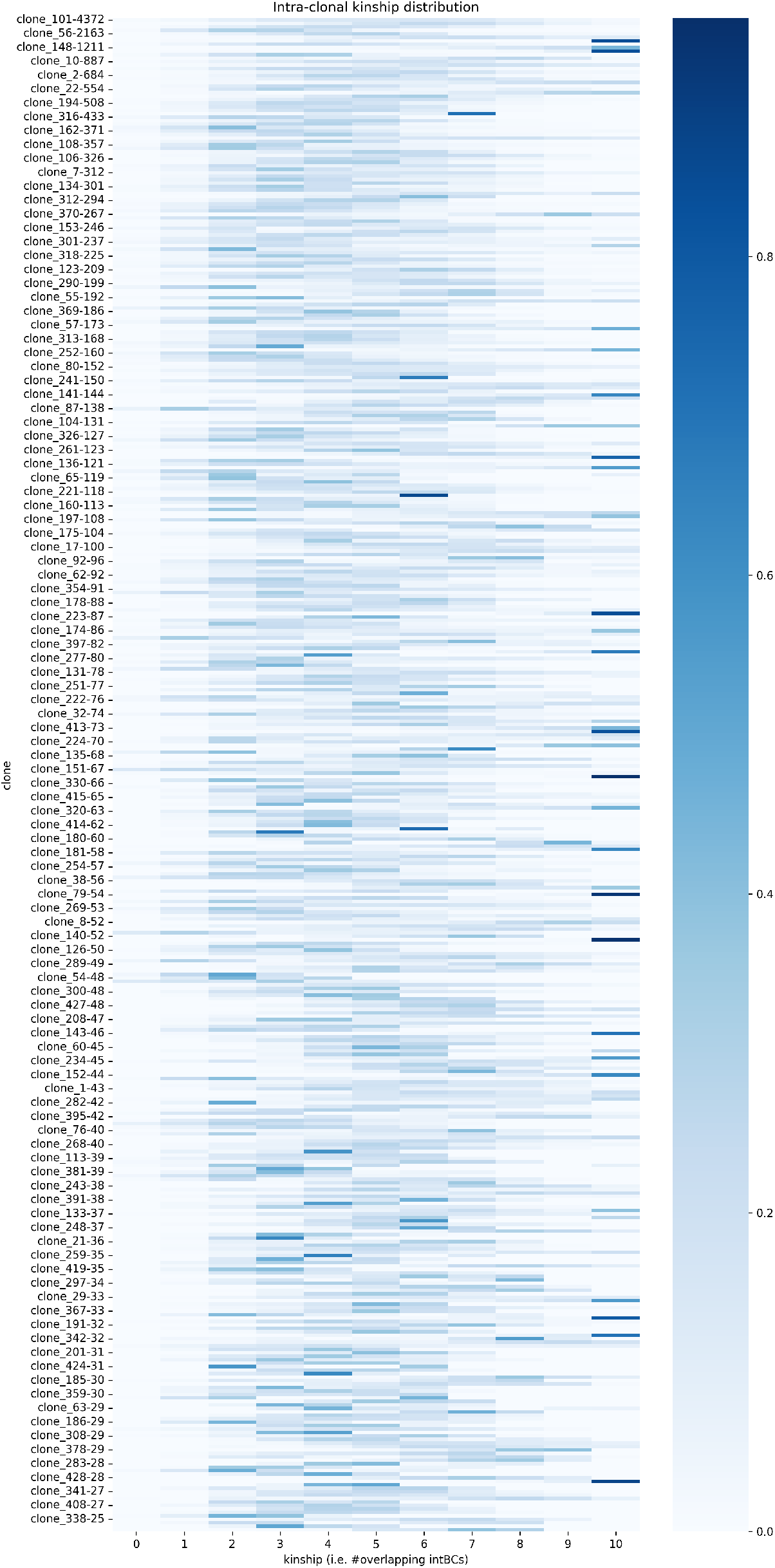
Quality control NovaClone pipeline output: Intra-clonal kinship. Example taken from real data. Each row corresponds to a clone, represented by its name and cell number. The kinship distribution of that clone is being shown (the kinship is capped at a max of 10). Each row adds up to exactly 1.

### 3.5 Inter-clonal kinship: homoplastic intBCs

The previous plot on intra-clonal kinship certifies that the cells within each called clonal population are coherent, but it does not certify that the cells across different clones are clonally *un*related. For example, if one takes a true clonal population and randomly splits it into two subsets, all the metrics discussed so far may look convincing, but in fact these two called clones should have been only one. Therefore, we can look at the *inter-clonal kinship*.

For this, we can simply look at the number of `intBCs` shared between the `intBC` sets of two clones. For example, if clone 1’s `intBC` set is {*AAA, BBB, CCC*} and clone 2’s `intBC` set is {*CCC, DDD, EEE, FFF, GGG*} then the inter-clonal kinship is 1. In most experimental designs, the random insertion of `intBCs` means that it is highly unlikely that two clones share more than one `intBC`. Therefore, we can check this in the clone call and we report it to the user.

### 3.6 Data retention

So far, the QC we have considered validates that the clonal populations are coherent and that pairs of clones are unrelated. However, it is still possible that the clone call has discarded a substantial portion of the `intBC` data while performing the clone call. For example, if a single clonal population is characterized by the `intBCs` {*CCC, DDD, EEE, FFF, GGG*}, the clone calling algorithm could just randomly distribute the cells into two groups, and in group 1 retain only `intBCs` {*CCC, DDD*} and in group 2 retain only `intBCs` {*EEE, FFF, GGG*} . In this case, the two groups will form clonally coherent clonal populations (i.e. with high intra-clonal kinship) which are also distinct (i.e. with low inter-clonal kinship). However, around 50% of the raw data has been discarded by the algorithm. Therefore, we turn to the raw data to certify that the algorithm has not discarded large amounts of data.

Specifically, we count how many molecules (i.e. `UMI_count`) in the raw data have been retained by the clone calling algorithm. For example, if in the raw data a cell has the following `UMI_count`: [(*AAA*, 50), (*AAB*, 6), (*BBB*, 30), (*CCC*, 23), (*ZZZ*, 2)] and the clone call specifies that this cell belongs to a clone with `intBC` set {*AAA, BBB, CCC*}, then 50 + 30 + 23 = 103 UMIs have been retained out of 111. What if the cell is a multiplet or has not been mapped to any clone? We can (1) choose to penalize the algorithm for this and consider this data as discarded, or (2) just limit ourselves to the successfully mapped cells - after all, we already have a QC metric that certifies that a majority of cells have been successfully mapped and the multiplet rate is coherent. We choose (2) in this case. Hence, we report, for the successfully mapped cells, how many of their `UMI_count` were retained by the algorithm. Moreover, we can plot the histogram of the distribution over all cells, as seen in Figure 10.

**Fig. 10.**
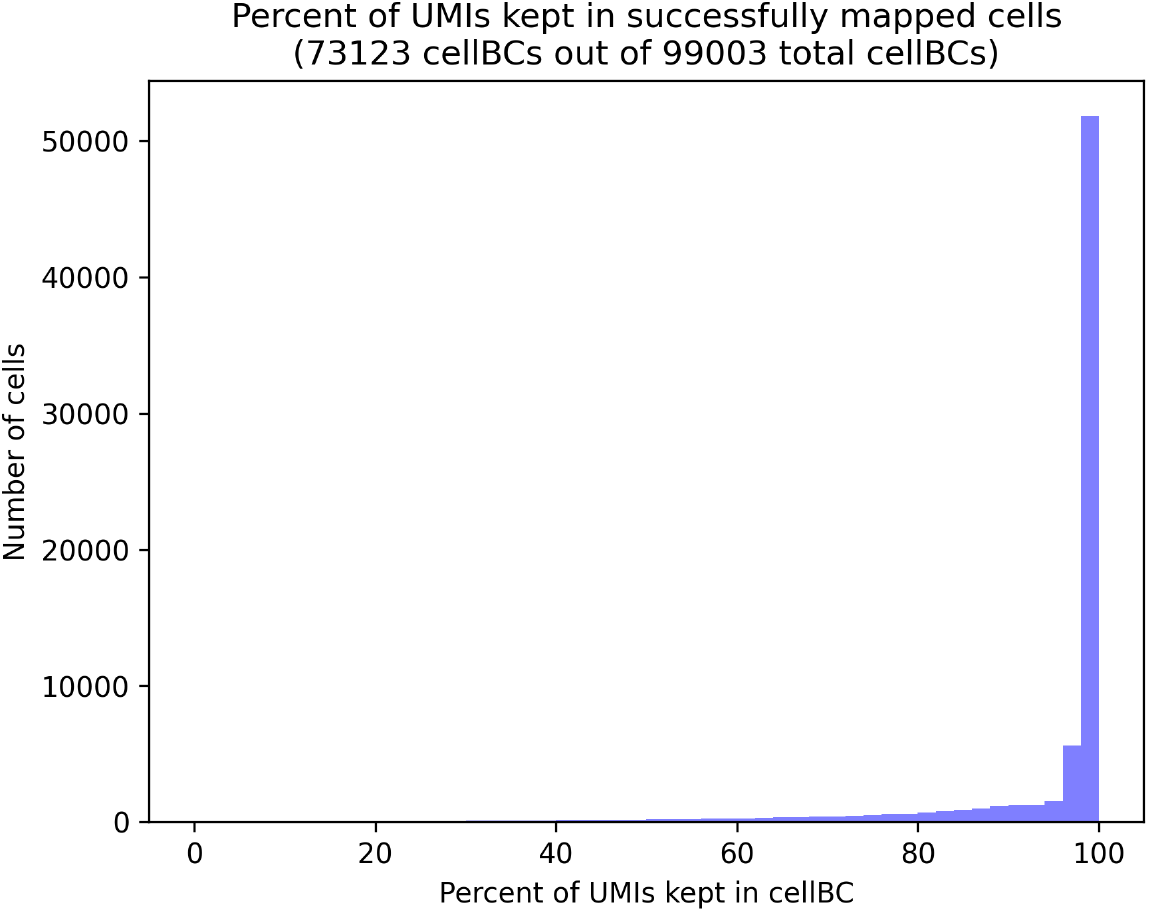
Quality control NovaClone pipeline output: Data retention. Example taken from real data. The x-axis shows the percentage of UMIs retained per `cellBC`, calculated as UMIs from `intBCs` passing the UMI filter divided by the total UMIs from all `intBCs` in the cell. The y-axis shows the number of cells. A distribution concentrated near 100% indicates that most UMIs are retained for downstream analysis.

### 3.7 Clone intBC sets

To assess the quality of the final `intBCs` clonal signature of each clone, we can inspect the frequency of each `intBC` in the *raw* data of that clone. By looking at the raw data, we include sequencing errors, homoplastic `intBCs`, background RNA – *anything* in the raw data will be considered. This gives a more critical view into what was included and *excluded* from the signature of that clone. We color-code `intBCs` in blue if they are included in the final clonal signature of the clone, and red otherwise. The `intBCs` are sorted in descending order of frequency.

We include additional information for each `intBC`: (i) the median `UMI_count` for that `intBC`, and (ii) the minimum edit distance to a more frequent `intBC` in that clone. This serves to reveal if any high-frequency `intBCs` were excluded due to being sequencing errors, background RNA, or other technical reasons. Figure 11 shows this plot for one clone in a real dataset. We show the top 20 `intBCs` though additional `intBCs` can be displayed as needed. In this figure, the data show that the clone has six high frequency `intBCs`. The lower-frequency `intBCs` are likely subclonal integrations (like *CCC* and *DDD* in the toy example from Figure 1). Indeed, the recursive clone call reveals significant subclonal structure for this clone, as seen in Supplementary Figure S1.

**Fig. 11.**
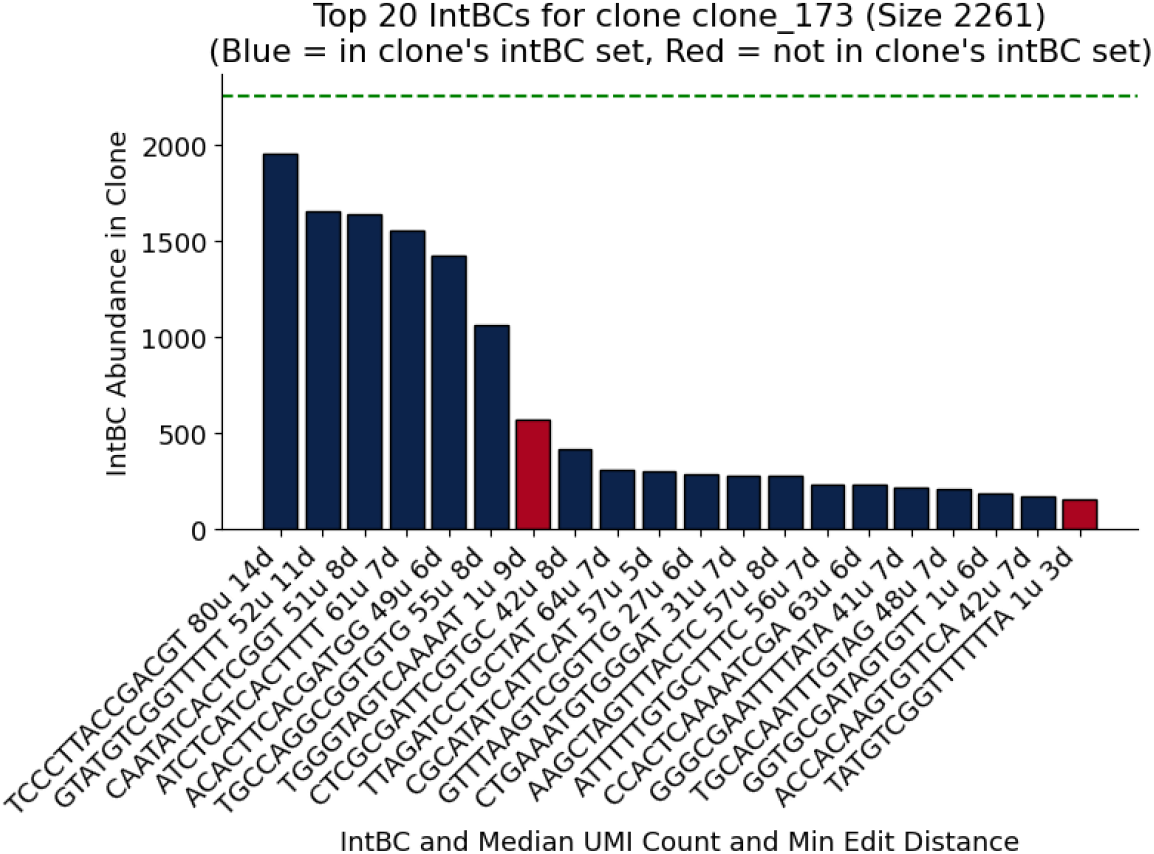
Quality control NovaClone pipeline output: Clone intBC sets. Bar plot showing the top 20 abundant `intBCs` in clone_173. Bars in blue represent `intBCs` from the clonal signature, while red bars represent `intBCs` present in the clone but not part of the clonal signature. The green line indicates the total number of cells in the clone (2,261). X-axis labels display the `intBC` sequence, mean `UMI_count`, and minimum edit distance from the nearest other `intBC`. In this example, the first red `intBC` (TGGGTA…) has a median `UMI_count` of 1 and a high edit distance to any more frequency `intBC`, suggesting it is background RNA. The last red `intBC` (TATGTC…) has a median `UMI_count` of 1 and a low edit distance of 3. It is most certainly a sequencing error of the second `intBC` (GTATGT…).

## 4 Benchmark

The performance of `NovaClone` was evaluated through a two-part benchmarking framework: **Part I: Synthetic Validation**. We first assessed the algorithm using simulated datasets that model the full lineage tree, from simulation of the evolution of lone lineages through clonal expansion, while incorporating various sources of technical noise. These simulations provided a ground truth to systematically compare `NovaClone` against re-implemented baseline approaches and established tools, across multiple performance metrics. The simulation parameters governing both topology and lineage barcoding are detailed in Table 1, Table **??**, Table 3. **Part II: Empirical Validation**. We further evaluated the algorithm’s utility on a publicly available dataset.

**Table 1.** Simulation parameters for clone lineage evolution.

| Symbol | Code Variable | Description | Default |
| --- | --- | --- | --- |
| $G$ | depth_tree_list | Actual depth of a specific tree, sampled from $[G_{\min}, G_{\max}]$. | [3, 5] |
| $G_{\max}$ | tree_depth_range_max | Maximum possible tree depth used to define barcode capacity. | 5 |
| $B_{\min\_gen}$ | min_barcodes_per_gen | Base capacity used to calculate total potential barcodes. | 10 |
| $\delta$ | decay_speed | Controls the steepness of integration rate change. | 1 |
| $\alpha$ | alpha | Trend of integration frequency (positive: increasing, negative: decreasing). | -1 |
| $P_{\min}, P_{\max}$ | minP, maxP | Operational bounds for individual integration probability. | 0.01, 0.9 |
| $\phi_{\text{total}}$ | perc_silenced_total | Target cumulative proportion of barcodes silenced. | 0.1 |
| $L$ | size_of_barcodes | Fixed length of the DNA barcode sequence. | 14 |

**Table 2.** Simulation parameters for clonal expansion and technical noise.

| Symbol | Code Variable | Description | Default |
| --- | --- | --- | --- |
| $(\mu_c, \sigma_c)_N$ | `lognorm_dist_cell_per_clone_params` | Parameters for log-normal clone size distribution. | Empirically derived |
| $N_{\min}$ | `min_cells_per_clone` | Minimum number of cells per terminal leaf. | 15 |
| $(\alpha_u, \beta_u, \text{loc}, \text{scale})_U$ | `beta_dist_UMIs_params` | Parameters for the UMI count distribution per barcode. | Empirically derived |
| $\epsilon$ | `error_rate` | Per-base probability of a substitution error. | 0.001 |
| $\gamma$ | `multiplet_rate` | Probability of cell-cell co-partitioning. | 0.05 |
| $\theta$ | `missing_data_rate` | Probability of an `intBC` dropout event. | 0.05 |

**Table 3.** Benchmarking parameter grid. Parameters were varied one at a time; all others were held at their default values. Each condition was run over five independent repetitions.

| Symbol | Default | Values tested | Description |
| --- | --- | --- | --- |
| $L$ | 14 | 4, 8, 10, 12, 14 | intBC sequence length |
| $\epsilon$ | 0.0001 | 0.0001, 0.001, 0.01, 0.05 | Per-base substitution error rate |
| $\theta$ | 0.05 | 0.01, 0.05, 0.1, 0.2, 0.3 | intBC dropout probability |
| $\gamma$ | 0.05 | 0.01, 0.05, 0.1, 0.2, 0.4 | Multiplet rate |
| $B_{\min\_gen}$ | 15 | 4, 7, 10, 15, 20 | Minimum barcodes per generation |
| $N_{\min}$ | 15 | 3, 5, 10, 15, 20 | Minimum cells per subclone |

### 4.1 Simulated dataset creation

#### Simulating the Evolution of Clone Lineages

We simulate a hierarchical tree structure representing clonal evolution over *G* generations, where *G* is randomly sampled from the range *G* ∈ [*G*_min_, *G*_max_] (Table 1). For each clone, we define the total barcode capacity as:

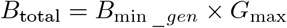

where *B*_min _*gen*_ represents the minimum number of barcodes per generation.

1. **Barcode integration probability** The probability of a barcode being integrated at any given generation *g* is determined by a non-linear scaling of the generation index. First, we calculate a normalized progress value (*x*_*g*_):

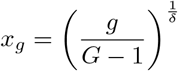

where *δ* is the decay speed. This value is used to determine *P*_*g*_, the generation-specific integration probability that is bound between *P*_min_ and *P*_max_ based on the trend parameter *α*:

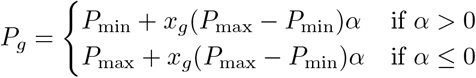
2. **Integration process** For every node in the tree, we perform a fixed number of integration trials *M*, where 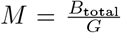. The integration of a barcode is determined by *M* independent trials. For each trial, an integration event occurs if a random draw from *U* (0, 1) is less than *P*_*g*_.
3. **Inheritance and epigenetic silencing** Upon cell division, daughter nodes inherit the set of `intBCs` from the parent. However, with a certain probability, an integration site can become inactive due to epigenetic silencing. We assume this state is heritable and remains inactive during all subsequent generations. To simulate heritable epigenetic silencing, we calculated a per-generation silencing rate *σ*_*g*_ such that the total percentage of `intBCs` lost across the entire tree matches the total silencing rate *ϕ*_total_:

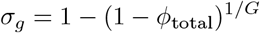

In each generation, every `intBC` has a probability *σ* of being silenced. Silenced `intBCs` are removed and not passed on to next generations.

At the end of this step, we obtain a simulated clone lineage tree, where the terminal nodes (leaves) represent subclonal progenitor cells, each with its corresponding set of `intBCs`.

#### Simulating Clonal Expansion and Noise

To generate realistic single cell datasets, we expanded the leaf nodes of the simulated lineage trees that represent subclonal progenitors into cellular populations and introduced various sources of technical noise.

1. **Clonal expansion and UMI assignment**. Each subclonal progenitor expands into a population of *N*_*c*_ cells. The population size of each subclone is sampled from a log-normal distribution (Table **??**) calibrated to represent the clonal size distributions observed in experimental data:

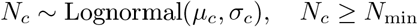

where *N*_min_ represents the minimum required subclone size. For each cell in the resulting population, every `intBC` is assigned a UMI count *U* . These counts are sampled from a scaled Beta distribution:

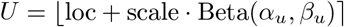

where the parameters (*α*_*u*_, *β*_*u*_, loc, scale) were calibrated to match experimental UMI distributions observed in droplet-based sequencing data.
2. **Technical noise sources**. Following expansion and UMI assignment, we simulate noise sources common to lineage tracing experiments:
  - **Substitution sequencing errors:** for every base in each `intBC`, a substitution error is introduced with a per-base probability *ϵ*. If an error occurred, the original nucleotide is replaced by one of the three alternative bases with equal probability.
  - **Multiplet formation:** To simulate technical artifacts we model cases where multiple cells are captured in a single droplet, causing them to be perceived as a single cells assigned the same `cellBC`. This is implemented by applying multiplet rate *γ*. Cells are randomly shuffled and grouped and a subset of these cells is then merged into a single “multiplet”, combining their `intBC` sets and `UMI_count` into a single `cellBC`. This process yield both intra-clonal and inter-clonal multiplets.
  - **Missing Data**: Dropout events are simulated by applying a probability *θ* to each `intBC` in the final counts table. If a dropout event occurs, the `UMI_count` for that `intBC` is set to zero, simulating cases where integration events are not detected.

The simulation pipeline concludes with the generation of a final `intBC` table containing `cellBCs`, `intBCs`, and `UMI_count`. This table incorporates all simulated technical noise and serves as the input for evaluating the clone calling performance. Notably, since clone calling operates solely on the `intBC` table, irrespective of whether barcodes were generated by static integration or dynamic CRISPR-based recording, this simulation framework is technology-agnostic and covers both experimental designs.

Following the generation of the synthetic datasets, we evaluated the baseline performance of the framework by executing `NovaClone` using its default parameter settings.

### 4.2 Public dataset analysis

To perform an analysis of experimental lineage tracing data, filtered dataset was obtained from *Simeonov et al*. [7]. This dataset was analyzed using `NovaClone` with default parameters except for threshold parameters that were set in the original analysis performed by *Simeonov et al*. of minimum edit distance = 2, minimum `UMI_count` per `intBC` = 2, and minimum clone size = 5 cells.

To compare the cell-to-clone assignments, we used the final classification provided in the original study. Silhouette scores were computed using scikit-learn [18] with the Jaccard similarity index as a metric, applied separately to the original and to `NovaClone` results using all clonal assignment of cells from each method.

For more direct comparison of cell-to-clone assignments, the original classification was filtered to include only cells with at least two unique `intBCs` and clones containing at least five `cellBCs`.

### 4.3 Baseline Clone Calling Methods

We implemented two commonly used strategies for clone calling as baseline comparisons: one based on Jaccard similarity with hierarchical clustering [4, 7, 19], and another using Pearson correlation of expression levels (`UMI_count`) followed by HDBSCAN clustering [20]. In both cases, input data was filtered similarly to the filtering in `NovaClone`, by both `UMI_count` per `intBC` per `cellBC` and by a minimum number of `cellBCs` expressing each `intBC`.

#### Jaccard and Hierarchical Clustering

After filtering using parameters `umi_threshold` (default=10) and `min_cells_with_intBC` (default=2), this method constructs a `UMI_count` matrix of cells and `intBCs`. Pairwise cell-to-cell Jaccard similarity is then calculated between `intBCs`, and the resulting distances matrix is used to perform hierarchical clustering with the method specified by parameter method_linkage (default=complete). The threshold for cutting the dendrogram is set to the maximum value of the dendrogram multiplied by a ratio that can be adjusted with parameter `ratio_dendrogram` (default=0.9). Clones are then filtered based on minimum number of `intBCs` per clone using parameter `min_intbc_per_clone` (default=2). Subsequently, `cellBCs` are assigned to clones based on the highest overlapping `intBCs`. Finally, clones are filtered based on parameter `min_cells_per_clone` (default=10).

#### Pearson and HDBSCAN Clustering

After filtering using parameters `min_UMIs` (default=10) and `min_cells_with_intBC` (default=10), this method constructs a `UMI_count` matrix of cells and `intBCs` and then pairwise cell-to-cell Pearson correlation between `intBCs` was computed. We then applied HDBSCAN clustering with key parameters `min_cluster_size` is `min_intbc_per_clone` (default=2) and `min_samples` (default=30) tuned for best performance on simulated data. Cells not assigned to any cluster by HDBSCAN were removed. Then, cells were assigned into the clone according to highest overlapping `intBCs`. Clones were filtered according to parameter of `min_cell_cluster_size` (default=30).

#### Cassiopeia clone calling Algorithm

After filtering the input data using the parameter `min_intbc_umi_count`, clonal populations were identified using the `call_lineage_groups` tool from Cassiopeia preprocessing pipeline using its default parameters [21]. This function identifies clones through an iterative kinship-based assignment, where cells are clustered based on the proportion of shared `intBCs`. The original output was transformed to match our benchmarking format.

#### One large clone

This model was generated by assigning all detected `cellBCs` to a single clonal lineage. It serves as a reference baseline for maximal over-clustering.

#### Empty clone call

This model was generated by not assigning `cellBCs` to any clonal lineage. It serves as a zero baseline where the algorithm returns an empty output.

### 4.4 Benchmarking Clone Calling

To evaluate the performance of the clone calling algorithms, we conducted benchmarking across multiple levels: multiplets detection, cell-to-clone assignment, and `intBCs` recovery. These assessments were performed both at the clonal level and at the subclonal level, which includes all possible nested clonal structures.

1. **Multiplets identification** was benchmarked by comparing predicted multiplets against the true set of simulated multiplets. Classification metrics including precision, recall, F1 score, and accuracy were computed. In addition, Jaccard similarity was used to assess the overlap between predicted and ground-truth multiplets sets.
2. **Cell-to-clone assignment** was evaluated by pairing each ground-truth clone with the most similar predicted clone, based on Jaccard similarity of their `cellBC` sets. For each matched clone pair, standard classification metrics (precision, recall, F1 score, accuracy) and the adjusted Rand index (ARI) were calculated.
3. `intBC` **recognition** was assessed within the matched clone pairs identified in the previous step. The sets of `intBCs` were compared using the same metrics: precision, recall, F1 score, accuracy, Jaccard similarity, and ARI. These metrics were also calculated in a weighted manner, where each `intBC`’s contribution is scaled by the number of cells in which it is detected.

Finally, we benchmarked subclonal recovery by calculating the proportion of `cellBCs` assigned to a subclonal lineage. This metric serves as a measure of **assignment completeness**, quantifying the fraction of the clonal architecture captured by the tool without assessing the topological correctness of those assignments.

## 5 Results

### 5.1 Benchmarking on Simulated Data

To evaluate the accuracy of `NovaClone`, we generated synthetic datasets that simulate the integration of `intBCs` process of lineage tracing experiments. The datasets include realistic `UMI_count` of `intBCs` per `cellBC`. We also simulated controlled levels of noise, including substitution sequencing errors, missing data, `intBC` sequence length variability, and varying multiplet rates to reflect common challenges in lineage tracing experiment data.

We compared `NovaClone` to three alternatives: Cassiopeia’s clone calling algorithm and two baseline methods, one based on Jaccard similarity followed by hierarchical clustering and one based on Pearson correlation and HDBSCAN clustering (Figure S2, Figure S3, Figure S4, Figure S5, Figure S6, Figure S7, Figure S8, Figure S9). Since each method defines clones at different resolutions, we benchmarked against the best matching subclonal structure for each method.

A key advantage of `NovaClone` is its robustness to noise, particularly in multiplets detection, which is a common challenge in droplet-based lineage tracing experiments. Our ability to detect multiplets contributes to accurate cell-to-clone assignments, even as the multiplet rate increases. In contrast, the baseline and Cassiopeia clone calling methods showed a clear drop in performance under high multiplet conditions (Figure 12A). Additionally, unlike other methods that remove potential multiplets, `NovaClone` assigns them to multiple clones, allowing them to be identified.

**Fig. 12.**
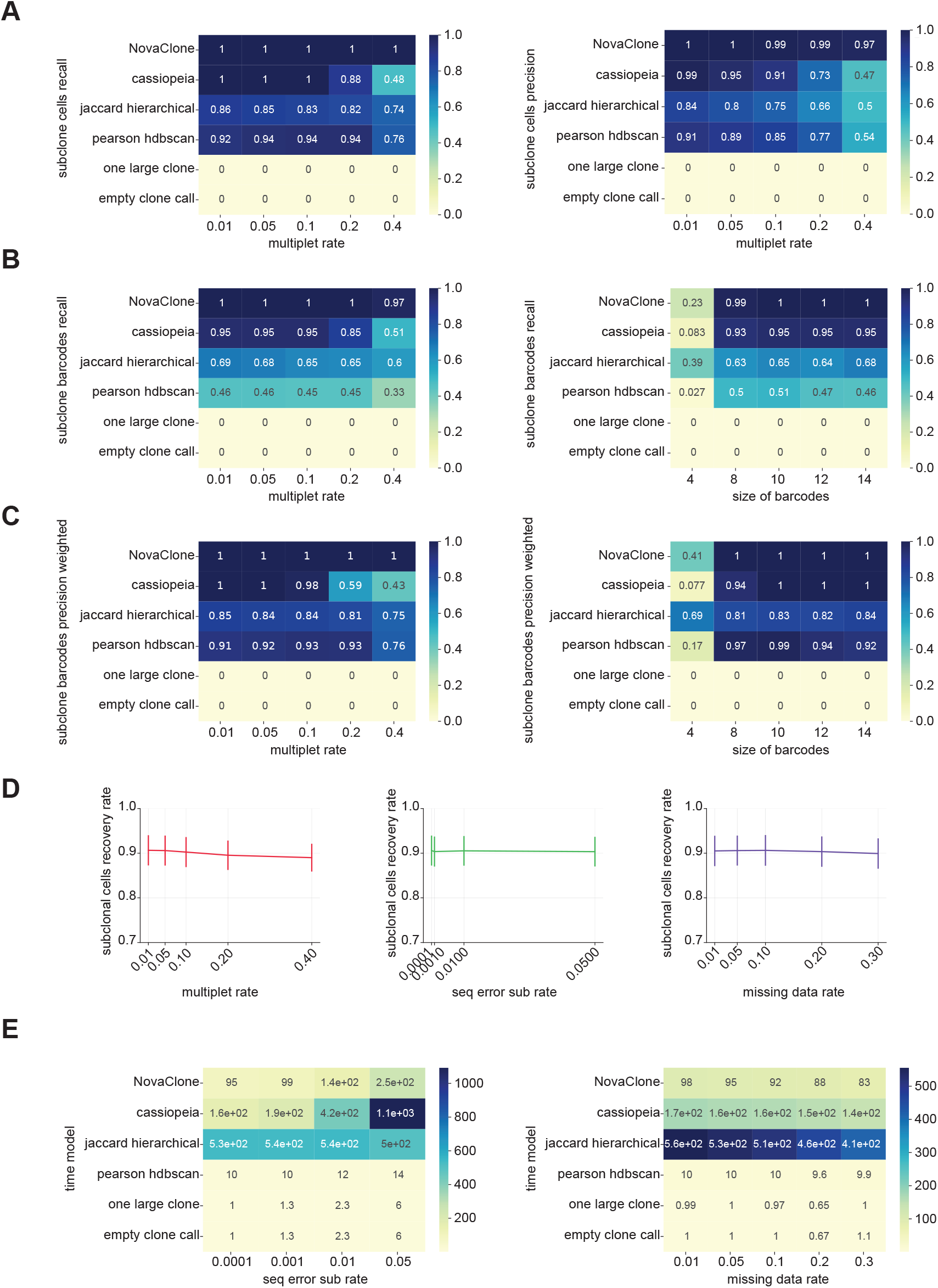
Benchmarking performance across simulated noise conditions. (A) Heatmaps showing cell-to-clone subclonal assignment performance across increasing multiplet rates. Left: recall values, right: precision values. (B) Subclonal `intBC` recognition performance measured by recall. Left: increasing multiplet rates, right: varying `intBC` lengths. (C) Subclonal `intBC` recognition performance measured by weighted precision. Left: increasing multiplet rates, right: varying `intBC` lengths. (D) Line plot showing subclonal `cellBC` recovery rates under different noise types: multiplet rate, sequencing error rate, missing data rate. (E) Runtime comparison of all methods under increasing noise. Left: substitution sequencing error rates, right: missing data rates.

In terms of `intBC` recognition, `NovaClone` achieved the highest recall across all noise conditions, effectively recovering the true `intBCs` while minimizing false negatives (Figure 12B). When weighting performance metrics by the number of cells expressing each `intBC`, `NovaClone` also showed high scores in precision and F1, highlighting its ability to both detect and correctly assign `intBCs` with high confidence (Figure 12C). While Cassiopeia showed similar performance of `intBC` recognition under high sequencing errors or missing data, `NovaClone` outperformed all alternatives in scenarios with high multiplet rates.

In addition to clone identification, `NovaClone` demonstrates a high subclonal assignment rate, successfully incorporating ~ 90% of cells into the subclonal structure. Notably, this assignment remains robust across technical noise, including sequencing error rates, missing data and increasing multiplet rates (Figure 12D).

These results showed that `NovaClone` can reliably identify both clones and subclonal structures, even under noisy conditions like high sequencing error rates, missing data, or high multiplet rates. On top of that, the short runtime makes it practical for larger datasets (Figure 12E). Overall, `NovaClone` offers both high accuracy and efficiency, which makes it useful for lineage tracing data processing.

### 5.2 Comparison to Published Experimental Data

To evaluate the performance of `NovaClone` on experimental data, we applied `NovaClone` to the *macs-GESTALT* static-`intBC` lineage tracing dataset from *Simeonov et al*. [7], which profiled clones in a metastatic pancreatic cancer model. As no ground-truth clonal labels exist for real lineage tracing data, we used the original study assignment as a reference to assess clustering quality. We compared the Silhouette scores of our clonal assignments with the clonal assignment from the original study. Using all cells assigned by each method, `NovaClone` achieved a higher Silhouette score (0.85 vs. 0.67; Figure 13A left), indicating more defined and separated clonal structures.

**Fig. 13.**
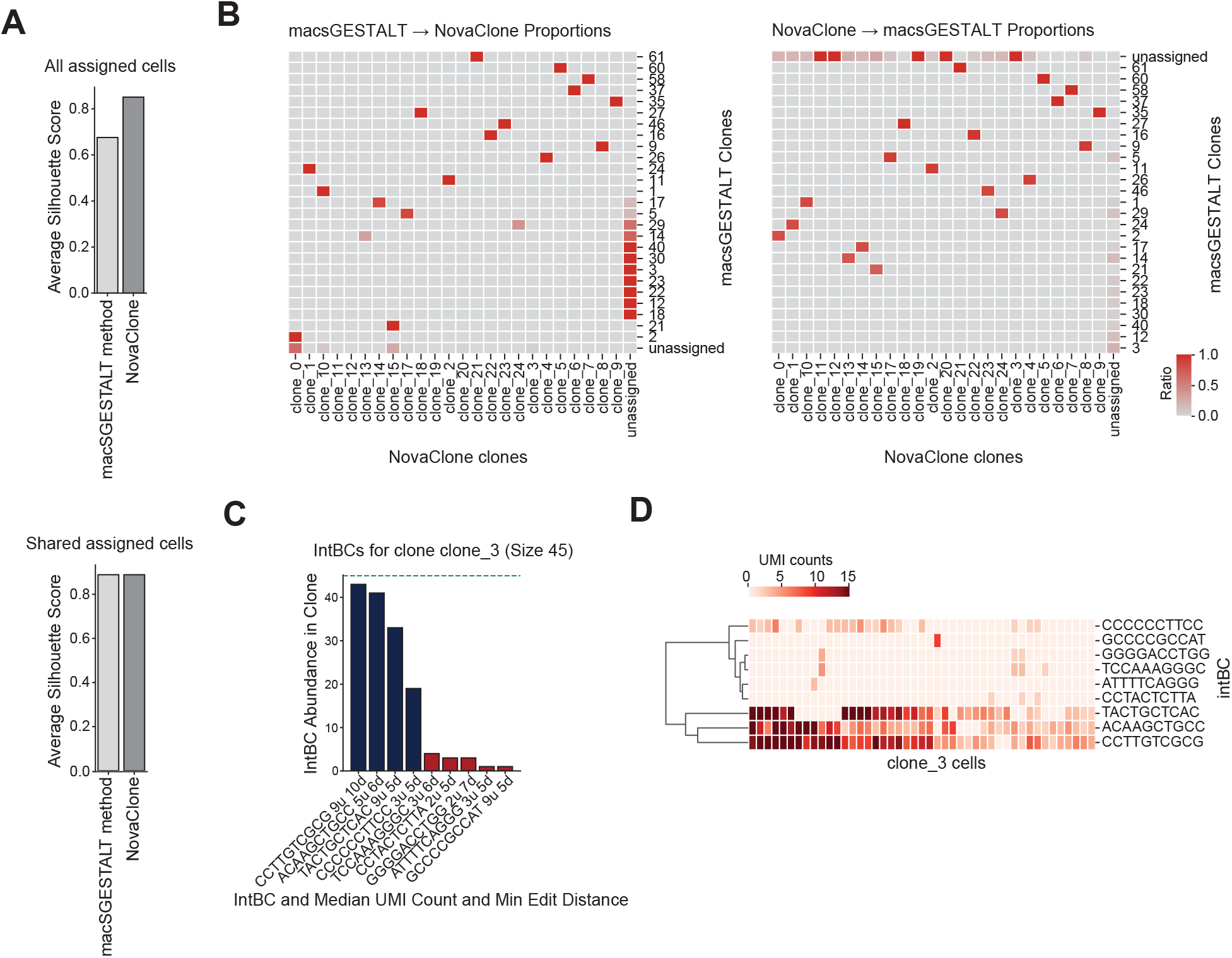
Comparison to published lineage tracing data. (A) Bar plot showing the average Silhouette scores for clones identified by the original *macsGESTALT* method and by `NovaClone`. Top: Based on all assigned `cellBCs`. Bottom: Based on shared assigned `cellBCs`. (B) Heatmaps showing the overlap in clonal assignments between the two methods. Left: proportion of `cellBCs` from each *macsGESTALT* clone found in `NovaClone` clones. Right: proportion of `cellBCs` from each of `NovaClone` clones found in *macsGESTALT* clones. (C) Bar plot showing the abundance of `intBCs` in clone_3. Bars in blue represent `intBCs` used as the clonal signature and red bars represent `intBCs` found in the clone but not used in the clonal signature. The green line indicates the total number of `cellBCs` in the clone. X-axis labels display the `intBC` sequence, mean `UMI_count` and minimum edit distance from the other nearest `intBC`. (D) Heatmap of `UMI_count` per `cellBC` for each `intBC` detected in clone_3.

To account for differences in data processing between the two methods, we performed an additional comparison using identical filtering constraints. Specifically, we restricted both analyses to a minimum of 2 `UMI_count` per `intBC` and a minimum clone size of 5 `cellBCs`. Under these conditions, both methods mapped 9,625 `cellBCs` to 25 clones with median clone size of 14 `cellBCs` (Figure S11). Since `NovaClone` requires at least two unique `intBCs` to define a reliable clonal signature, we applied comparable filtering to the published results by excluding `cellBCs` with less than two unique `intBCs` and retaining only clones with at least 5 `cellBCs`. After applying comparable filtering to both methods, we reassessed clustering quality based on the intersection of assigned `cellBCs` (n=7,721), both methods obtained a Silhouette score of 0.88, demonstrating that `NovaClone` provides consistent and high-quality clustering that is fully aligned with the published results when using uniform filtering parameters (Figure 13A right).

We then examined the clones identified in the original study that were not recovered by `NovaClone`. Among those 7 clones, all either contained only one dominant `intBC` or lacked a consistent `intBC` signature across `cellBCs`. For example, clones 3 and 23 each had one dominant `intBC`, which is unreliable for confident clone calling and could instead result from background expression or contamination. Furthermore, in clone 3, the dominant `intBC` is also homoplastic, and in our results, it appears across three distinct clones. Similarly, clones 40 and 22 each have two `intBCs` with a small edit distance from one another and probably result from a sequencing error that reduces them to one dominant `intBC`, again making them unreliable clones. Other clones showed inconsistent `intBC` expression patterns. For instance, in clone 18 (36 cells), the two `intBCs` were each expressed in ∼ 20 `cellBCs`, but with minimal overlap, suggesting the absence of a coherent clonal signature (Figure 13B left, Figure 13A-B). In clones 14 and 29, only a subset of `cellBCs` overlapped with clones identified by `NovaClone`. In both cases, `intBC` co-expression was low, and `NovaClone` instead found smaller clones with better-defined and consistent clonal signatures (Figure S12C).

In contrast, `NovaClone` identified 5 clones (ranging from 5 to 45 cells) that were not assigned by the original method but showed strong clonal signatures. These clones had at least two `intBCs` consistently expressed across almost all the `cellBCs` in the clone, suggesting a reliable clonal signature. One illustrative example is clone_3, containing 45 `cellBCs` and 9 unique `intBCs`. Of these, 3 `intBCs` were present in nearly all `cellBCs`, 1 in ∼ 50% of `cellBCs`, and the remaining 5 in smaller subsets (Figure 13B right, Figure S13A-B). Also, the `UMI_count` for the clonal signature `intBCs` were high across the `cellBCs`, which further supports their reliability as a clone (Figure 13C-D).

Taken together, these results show that `NovaClone` not only recovers more well-defined clones, but also avoids assigning `cellBCs` to clones with weak or inconsistent clonal signatures, enabling more reliable clone calling for downstream analysis.

## 6 Discussion

In this work, we have introduced `NovaClone`, a principled and broadly applicable algorithm for identifying clonal populations in single-cell lineage tracing data, together with a comprehensive set of quality control diagnostics that can be applied to the output of any clone calling algorithm – not just our own. `NovaClone` is broadly applicable to lineage tracing technologies that rely on the integration of distinct `intBCs` into cells, whether sequential or static, making it compatible with a wide range of experimental designs.

`NovaClone` relies on a network-based algorithm, using concepts similar to those of other successful methods such as *UMITools*. At a high level, `NovaClone` works by building a graph of `intBC`-`intBC` co-occurrences, and uses statistical filters to extract robust clonal signatures (`intBC` sets) from the graph. These are then used to map cells to clones, call multiplets, and reconstruct the complete `intBC` sets, including lower frequency `intBCs` that were not a part of the clonal signature.

The only major assumption of our method is that two clonal populations must share at most 1 `intBC`. This is met, with overwhelmingly high probability, for any standard lineage tracing technology by virtue of the high diversity of the `intBC` space.

Using synthetic datasets, we demonstrate that `NovaClone` consistently outperforms existing clone calling approaches under multiple sources of noise, including high multiplet rates, substitution sequencing errors, missing `intBCs`, and varying `intBC` lengths. A key strength of `NovaClone` is its ability to detect multiplets, which helps maintain accurate cell-to-clone assignments even in noisy conditions. In addition, by effectively identifying the core `intBCs` of the clone, and allowing recursive analysis, our approach can resolve both clonal and subclonal structures at high resolution and accuracy. Beyond accuracy, `NovaClone` is also computationally efficient, allowing the analysis of large-scale datasets with minimal runtime.

We further evaluated `NovaClone` using the published *macsGESTALT* lineage tracing dataset from a metastatic pancreatic cancer model. `NovaClone` recovered many of the clones identified in the original study and additionally found well-defined clones, such as clone_3. Some of the clones reported in the original analysis were not recovered by `NovaClone`, mainly due to non-coherent `intBC` signatures or homoplastic `intBCs` that make confident clone identification unreliable. It is important to note that the original study, like many clone calling analyses of lineage tracing datasets, used a complex pipeline that included several manual or dataset-specific steps. In comparison, `NovaClone` requires minimal parameter tuning and can be easily applied to other datasets.

Taken together, `NovaClone` offers a practical approach for identifying clonal structures in single-cell lineage tracing data. As lineage tracing technologies continue to expand in areas such as developmental biology and cancer research, methods that can perform clone calling across diverse datasets and technologies with minimal tuning are becoming increasingly useful. `NovaClone` combines accuracy with computational efficiency, making it suitable for analyzing a wide range of lineage tracing experiments.

Our clone calling algorithm and QC metrics, together with code to seamlessly reproduce all results in this work, are available in the open source `nova-clone` Python package.

## 7 Conclusions

We introduce `NovaClone`, a network-based clone calling algorithm for single-cell lineage tracing datasets that is compatible with a range of experimental designs and technologies and requires minimal hyperparameter tuning. Benchmarking on synthetic datasets with realistic noise and on published data demonstrated high performance in clone assignment and `intBC` recognition. As lineage tracing becomes central to research in development, cancer evolution, and tissue dynamics, our scalable and accurate tool addresses a critical need for reliable analysis of large lineage-tracing datasets. We make our algorithm and QC metrics available through the open source `nova-clone` Python package.

## 8 Data Availability

The single-cell dataset analyzed in this study is publicly available and can be accessed through GEO under accession number GSE173958 and in Mendeley Data: https://doi.org/10.17632/t98pjcd7t6.1. All code used for analysis and simulation is available at: https://github.com/YosefLab/nova-clone. Simulated datasets were generated using this repository and can be reproduced using the simulation framework described in the Methods.

## 9 Supplementary Material

### 9.1 NovaClone parameters

Here we describe the parameters of `NovaClone`’s Python API. These can be grouped into three sets of parameters: (i) parameters for phase 1 of the algorithm, (ii) parameters for phase 2 of the algorithm, and (iii) parameters for phase 3 of the algorithm.

The parameters for the first phase of the algorithm (finding clonal signatures) are as follows:

- `intbc_table_per_sample`: Each value (`intBC` table) should be a pandas DataFrame with the following columns: `cellBC` (str), `intBC` (str), `UMI_count` (int). For example, a row might be: [“AAACCCAAGAGCCTGA”, “ACCTCATGATATTC”, 45] indicating that the `intBC “ACCTCATGATATTC”` was observed 45 times in the `cellBC “AAACCCAAGAGCCTGA”`. There should be one `intBC` table per sample; the sample name acts as the key of the dictionary.
- `min_intbc_umi_count` (default 10): `intBCs` with less than this number of `UMI_counts` will be discarded by the algorithm. The default value is 10, which is effective at removing sequencing errors and background RNA. For some datasets, such as the macsGESTALT which we analyze, we lower this threshold to two UMIs, as described in the main text. More generally, we recommend to set the UMI cutoff by considering a histogram of UMI counts across all `(cellBC, intBC)` pairs (rows in the input table) and use the leftmost mode (in case the count distribution is multi-modal) or a low quantile (otherwise).
- `custom_intbc_filtering_function` (default `lambda intbc: False`): A function taking in an `intBC` and returning `True` or `False`. `intBCs` for which this function gives `True` will be discarded. This is done in the step ‘Filtering spurious entries from the input table’. This is helpful when, for instance, it is known that `intBCs` should have a fixed length (e.g. 14). Of course, the user could in principle filter these `intBCs` out *before* calling our algorithm, but we strove to make our algorithm fully self-contained. By default, no filtering is performed.
- `min_intbc_cellbc_count` (default 10): `intBCs` appearing in less than this number of `cellBCs` will be discarded. This is done in the step ‘Filtering spurious entries from the input table’. This accomplishes the goal of removing rare `intBCs` which are thus less helpful for identifying clones. See the section ‘Pruning spurious edges’ for a discussion of the effectiveness of using a threshold of 10 (rather than a lower or higher one). We thus do not recommend lowering this threshold below 10.
- `edit_distance_cutoff` (default 4): Minimum *allowed* edit distance. For instance, if you expect there to exist sequencing errors at edit distance 3 but not 4, please set this to 4. This parameter is used as follows: When building clonal signatures (phase 1 of the algorithm), we identify putative sequencing errors by checking for `intBCs` *x* frequently co-occurring with another (more frequent) `intBC` *y* at an edit distance ≤ `edit_distance_cutoff`. If so, we remove *x*. In the second phase of the algorithm, when we reconstruct the `intBC` set for each clone de novo, we require that the edit distance to any other more abundant `intBC` is ≥ `edit_distance_cutoff`. Note that our cutoff for phase 1 is more stringent to be safe, since sequencing error in phase 1 have a bigger consequence (*over-merging* clones) than in phase 2 (just getting some aliased characters which will likely get dropped due to low frequency via `min_intbc_fraction` anyway). If desired, we allow setting the cutoff for phase 2 differently with `edit_distance_cutoff_phase_2`.
- `min_cells_assigned` (default 30): Coresets with less that this number of mapped cells will be dropped. If you wish to recover smaller clones, you can lower this, however, note that since all `intBCs` are supported by at least `min_intbc_cellbc_count cellBCs` (by default 10, and we do not recommend changing this), all clones will have at least 10 mapped `cellBCs`.

The parameters for the second phase of the algorithm (mapping `cellBCs` to clonal signatures and calling `intBC` sets de novo) are as follows:

- `min_intbc_fraction` (default 0.20): We only add `intBCs` to the clone’s (de novo computed) `intBC` set if it appears in at least this fraction of `cellBCs`. This filter is common in lineage tracing analysis as `intBCs` appearing in few cells are usually discarded. However, one can relax this threshold all the way down to zero to recover rare subclonal `intBCs` (e.g. those resulting from late integrations and subclonal structures). When using the recursive version of the algorithm, we require this to be set to zero (or else only shallow subclonal structures may be found). Please note that in addition to this fraction threshold, we require each `intBC` to appears in at least 10 cells in the clone, so even setting `min_intbc_fraction` = 0 will produce high-quality `intBC` sets.
- `edit_distance_cutoff_phase_2` (defaults to same value as `edit_distance_cutoff`): See description of parameter `edit_distance_cutoff`. If None (default) this is set equal to `edit_distance_cutoff`. This should work well in most cases. Reasons for setting this value to something different than `edit_distance_cutoff` are (i) increase retention of `intBCs` after clone calling; note that while removing sequencing errors during the first phase (finding clonal signatures) is critical (or else we might overmerge clones), when building `intBC` sets de novo is it less critical because in the worst case, sequencing errors lead to ‘duplicate’ characters which will likely get dropped due to low frequency via `min_intbc_fraction` anyway, and (ii) the user might want to curate the `intBCs` in each clone *themselves*, in which case they can set `edit_distance_cutoff_phase_2`=0 to turn off this second sequencing error filter.

The parameters for the recursive clone calling step are:

- `recursive` (default `False`): If `True`, then we will recursively call the algorithm on each clone, thus resolving the subclonal structure derived by the hierarchy of integrations. For this to work best and recover deep integration hierarchies, we require `min_intbc_fraction` = 0 (or else only shallow subclonal structures may be found).
- `recursive_min_fraction_uniquely_mapped_cells_to_recurse` (default 0.7): If recursive is True, then this determines the min fraction of cells that have to be uniquely mapped to a clone’s subclones before we accept the subclonal decomposition of that clone. This thus serves as a stopping criteria. For example, a clone might have two resolvable subclones comprising 35% and 31% of the clone’s cells respectively, with the remaining 34% unmapped (i.e. discarded). Hence, if this argument is set to 0.7 (i.e. 70%), then we would NOT accept this recursive decomposition and just retain the original clone. Instead, if the subclones comprised 35% and 35% of the clone’s cells, then we would exactly meet the 70% threshold and accept the recursive decomposition.
- `recursive_min_cells_assigned` (default 10): If `recursive` is `True`, a lower bound on the size of the subclones we want to find. Subclones tend to be smaller than the parent clone, so it is a good idea to set this lower than the original `coreset_dealiasing_step min_cells_assigned`. The default of 10 should work well.

Internal parameters of the algorithm, such as the 0.5 cutoff for identifying reliable edges, are exposed as private parameters with underscores in the API (e.g. `_reliable_edges_step ratio_cutoff`). We do not expect the user to ever have to change these, as they are robust across datasets based on our theoretical analysis (see main text); all the results in this paper were obtained with default values for all the private parameters.

### 9.2 Benchmarking Parameter Grid

We benchmarked `NovaClone` by varying one noise or structural parameter at a time, holding all other parameters fixed at their default values. Each condition was evaluated over five independent repetitions with different random seeds. Table 3 summarizes the parameters varied and the values tested.

### 9.3 Subclonal structure of clone 173

**Fig. S1.**
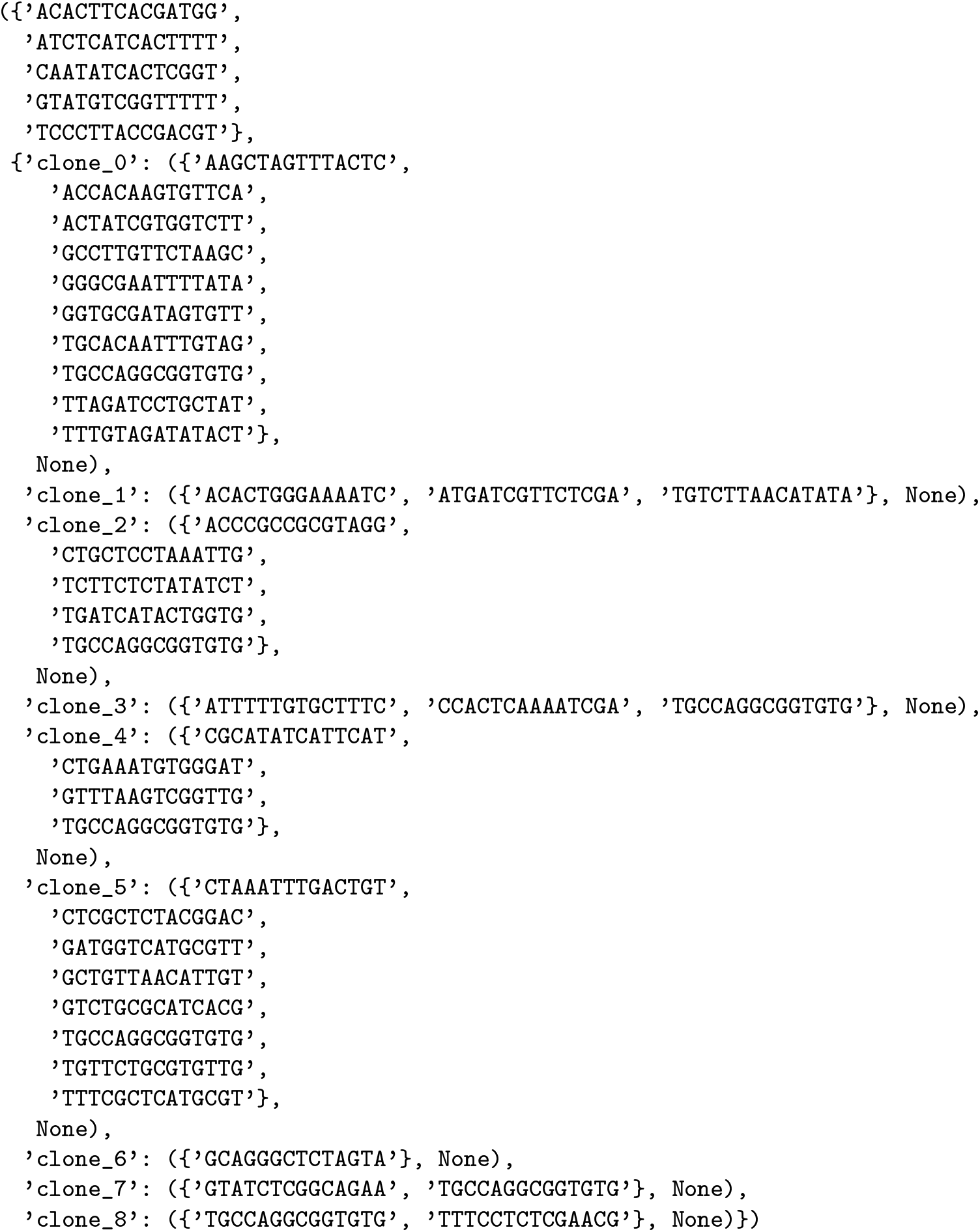
Subclonal structure on a real dataset. The subclonal structure of clone 173 found by our algorithm. The progenitor `intBC` set consists of the 5 `intBCs ACACTTCACGATGG`, ATCTCATCACTTTT, CAATATCACTCGGT, GTATGTCGGTTTTT, TCCCTTACCGACGT, which are shared by all cells in the clone. Thereafter, 9 subclones are identified. We used a fraction of *p*_accept_ = 50%.

**Fig. S2.**
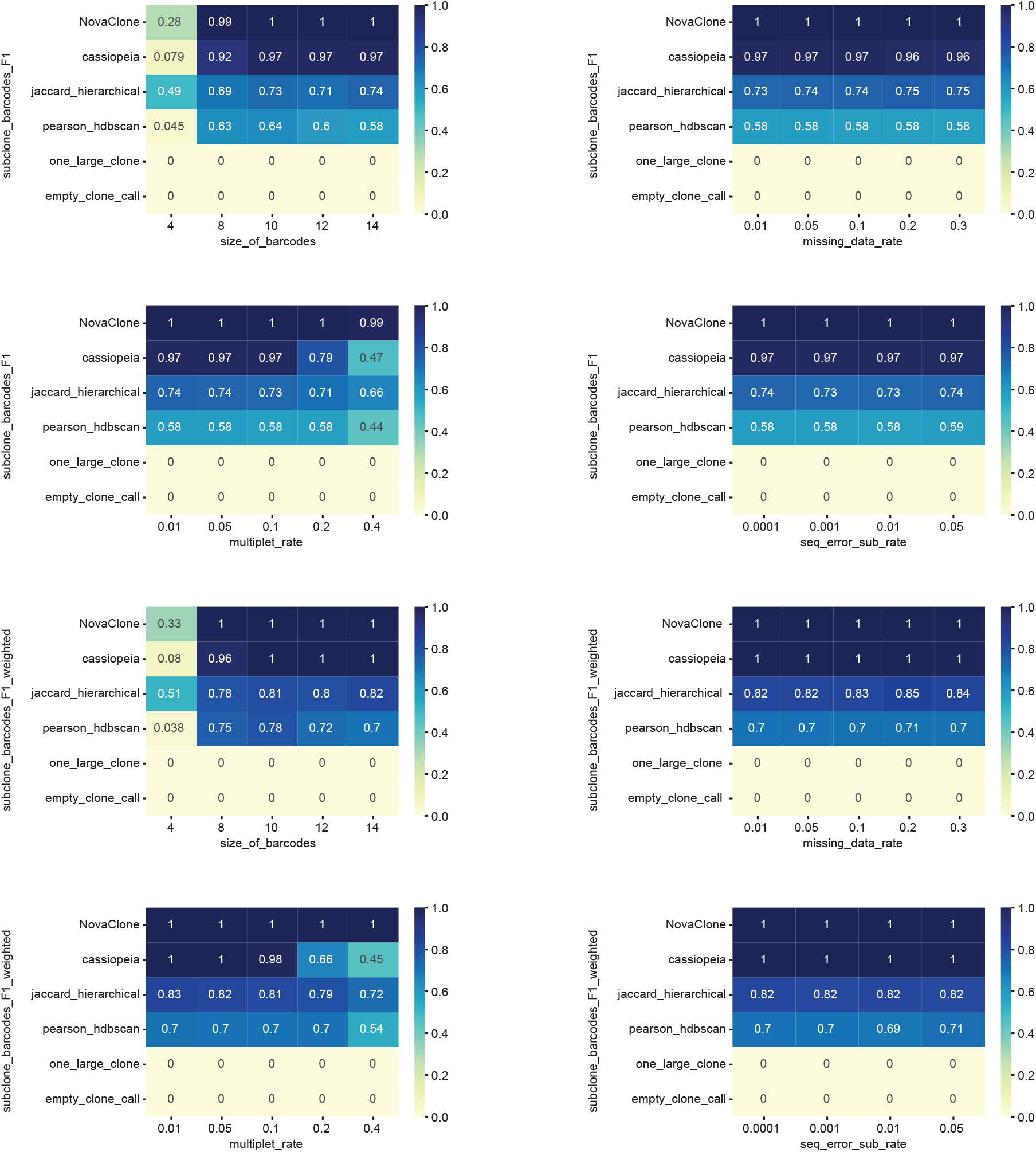
Benchmarking barcode F1 under simulated noise. Each heatmap shows F1 values across different noise levels for a given noise type. Rows correspond to methods and columns correspond to increasing levels of noise. Color indicates F1 values ranging from 0 to 1.

**Fig. S3.**
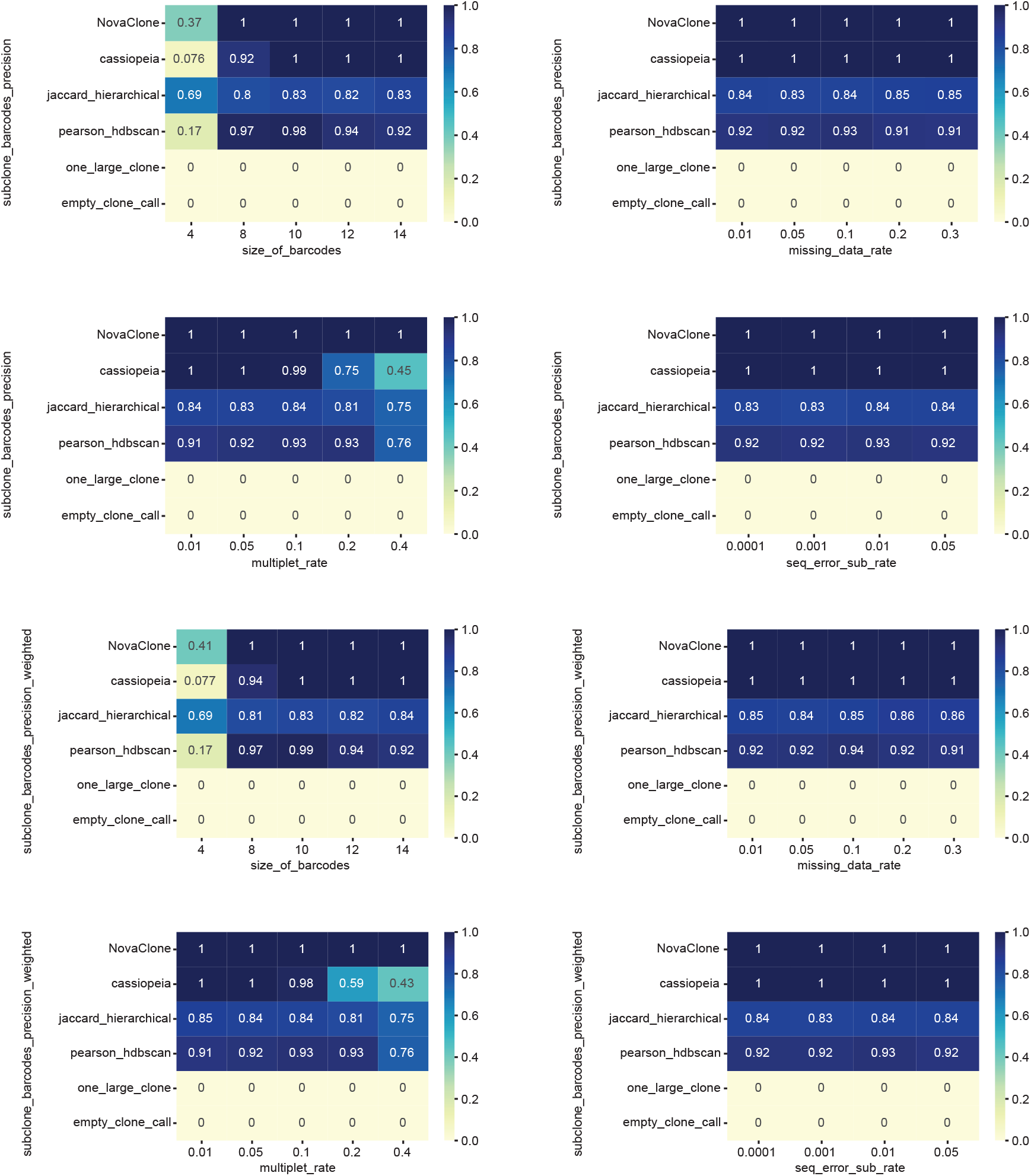
Benchmarking barcode precision under simulated noise. Each heatmap shows precision values across different noise levels for a given noise type. Rows correspond to methods and columns correspond to increasing levels of noise. Color indicates precision values ranging from 0 to 1.

**Fig. S4.**
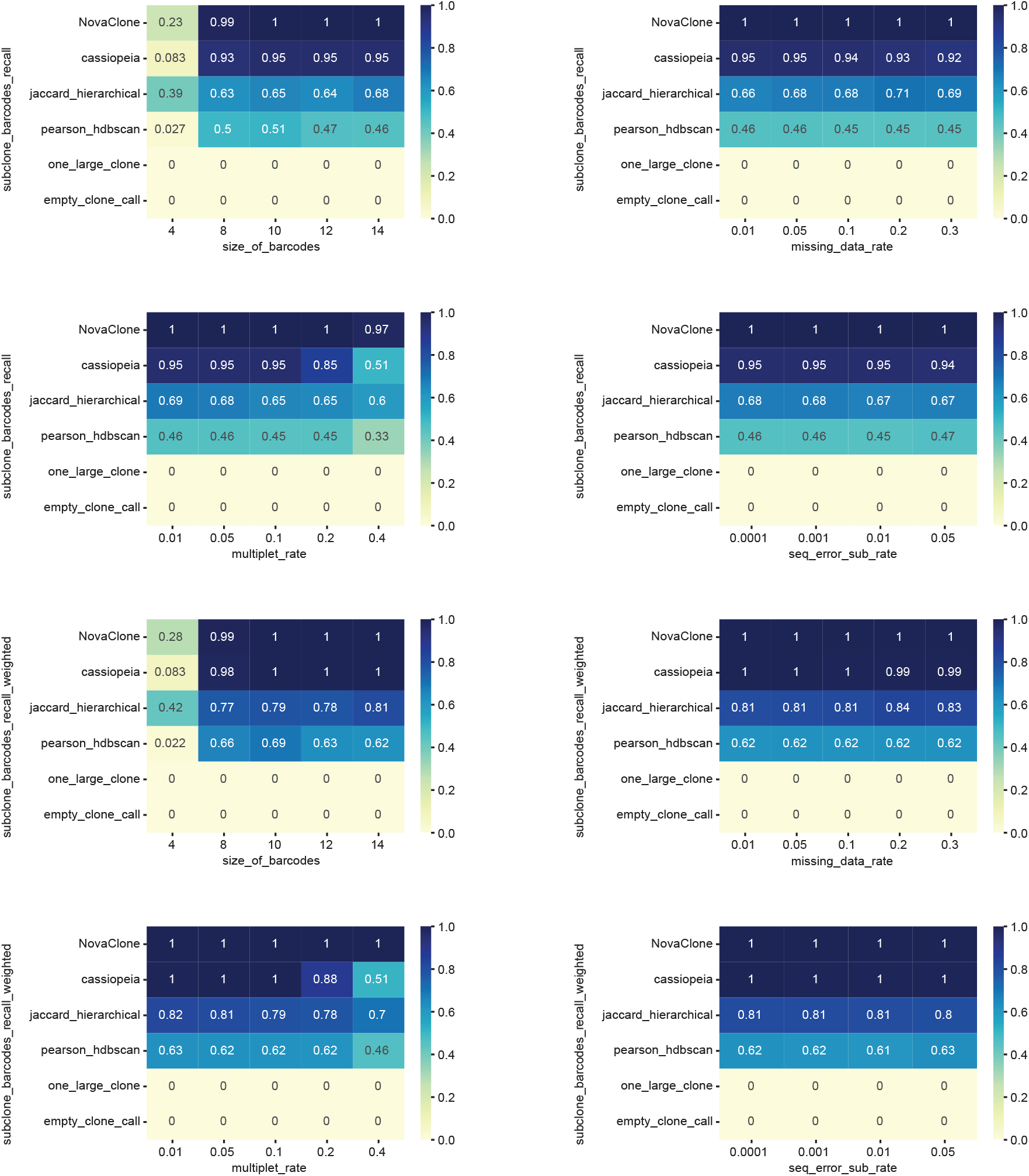
Benchmarking barcode recall under simulated noise. Each heatmap shows recall values across different noise levels for a given noise type. Rows correspond to methods and columns correspond to increasing levels of noise. Color indicates recall values ranging from 0 to 1.

**Fig. S5.**
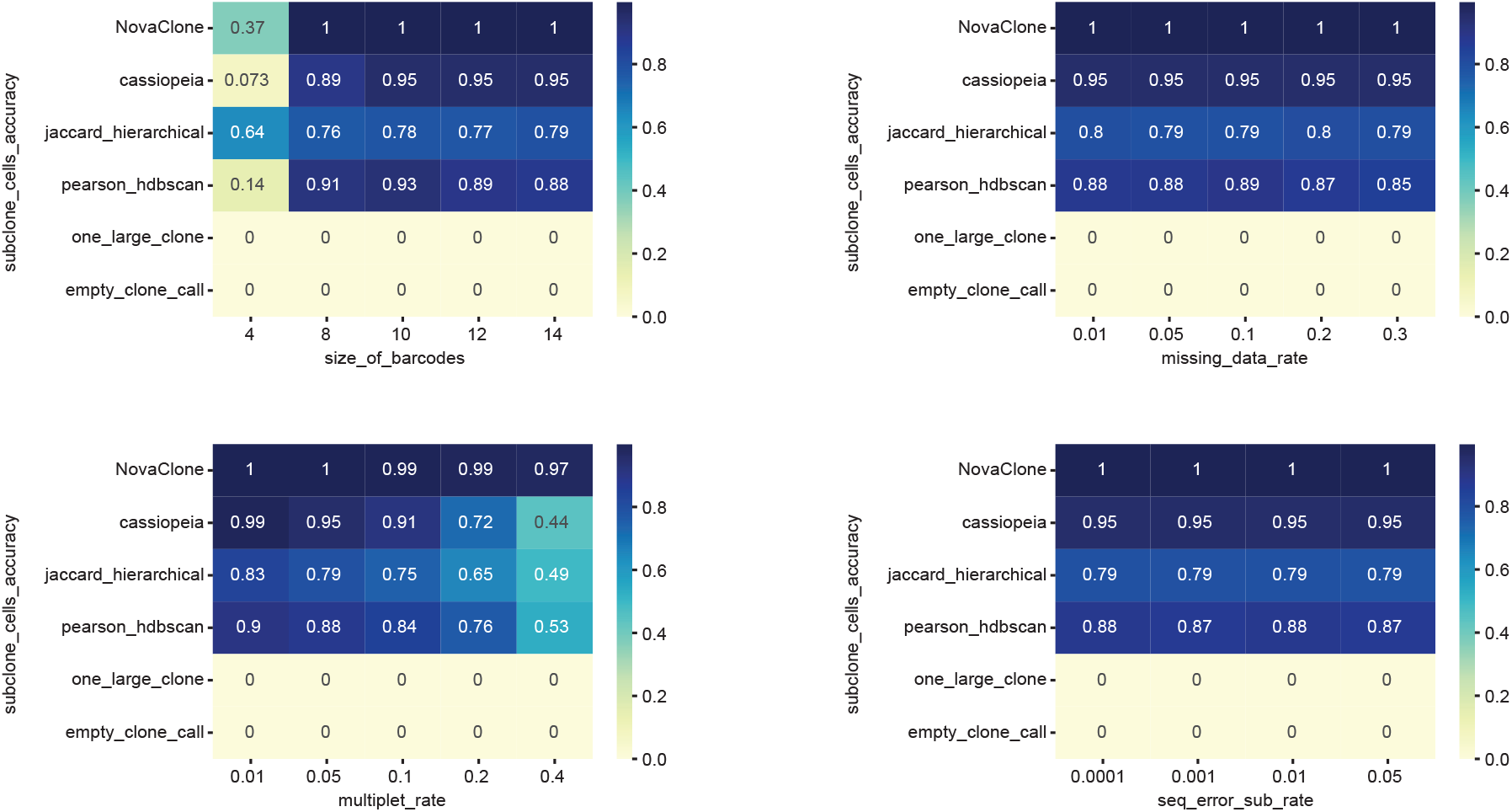
Benchmarking cell-to-clone accuracy under simulated noise. Each heatmap shows accuracy values across different noise levels for a given noise type. Rows correspond to methods and columns correspond to increasing levels of noise. Color indicates accuracy values ranging from 0 to 1.

**Fig. S6.**
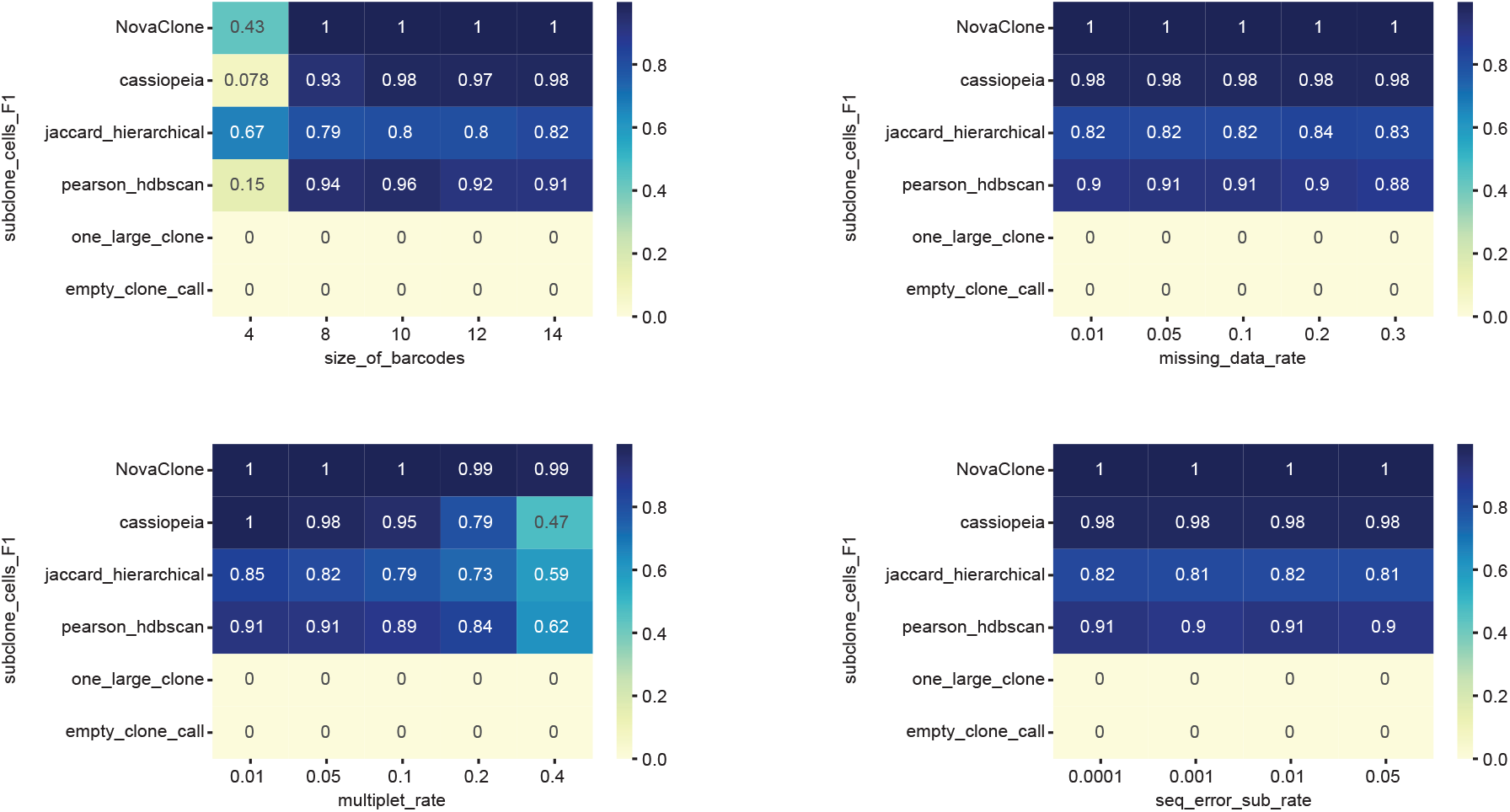
Benchmarking cell-to-clone F1 under simulated noise. Each heatmap shows F1 values across different noise levels for a given noise type. Rows correspond to methods and columns correspond to increasing levels of noise. Color indicates F1 values ranging from 0 to 1.

**Fig. S7.**
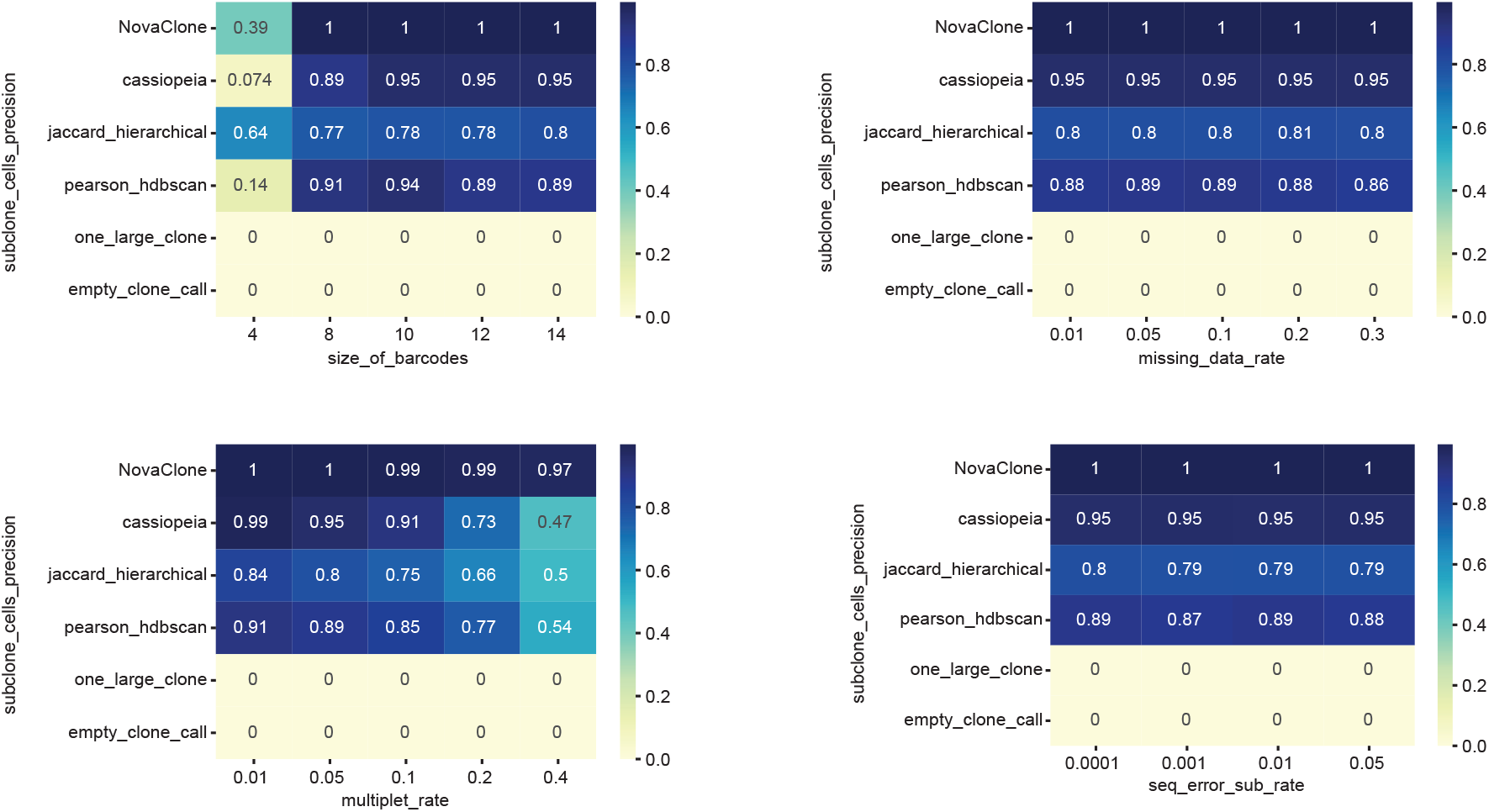
Benchmarking cell-to-clone precision under simulated noise. Each heatmap shows precision values across different noise levels for a given noise type. Rows correspond to methods and columns correspond to increasing levels of noise. Color indicates precision values ranging from 0 to 1.

**Fig. S8.**
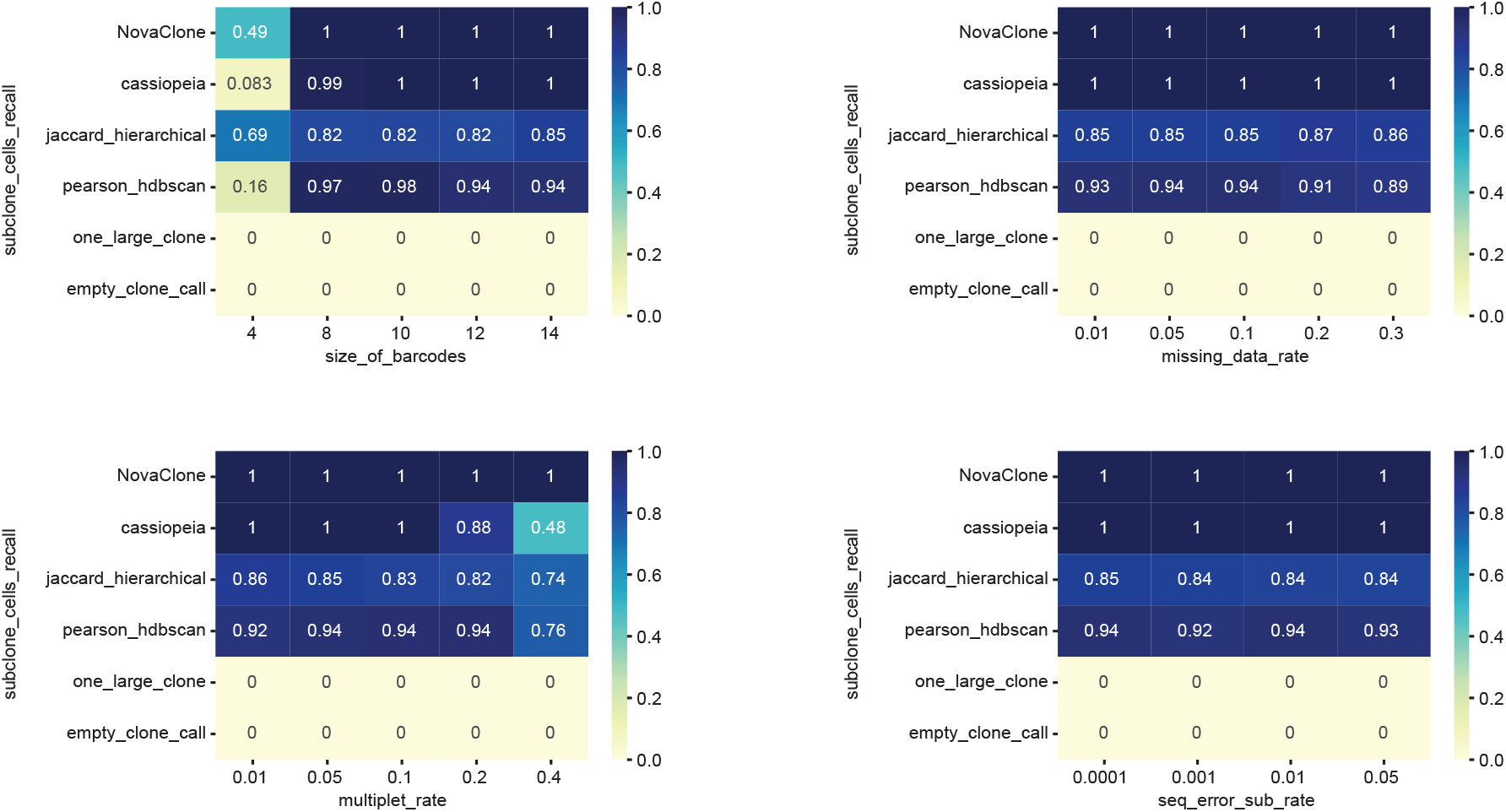
Benchmarking cell-to-clone recall under simulated noise. Each heatmap shows recall values across different noise levels for a given noise type. Rows correspond to methods and columns correspond to increasing levels of noise. Color indicates recall values ranging from 0 to 1.

**Fig. S9.**
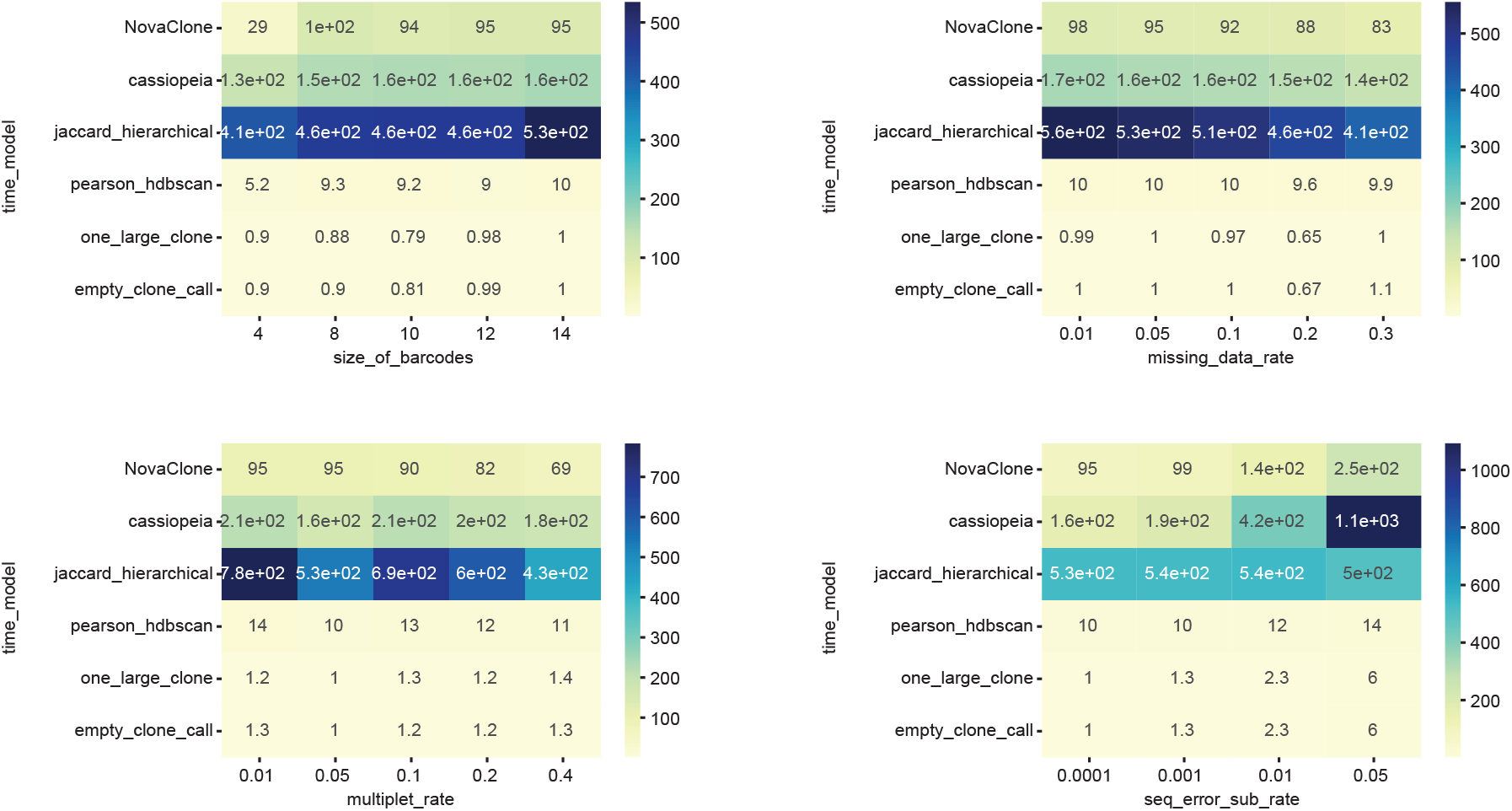
Benchmarking run time under simulated noise. Each heatmap shows run time across different noise levels for a given noise type. Rows correspond to methods and columns correspond to increasing levels of noise. Color indicates run time.

**Fig. S10.**
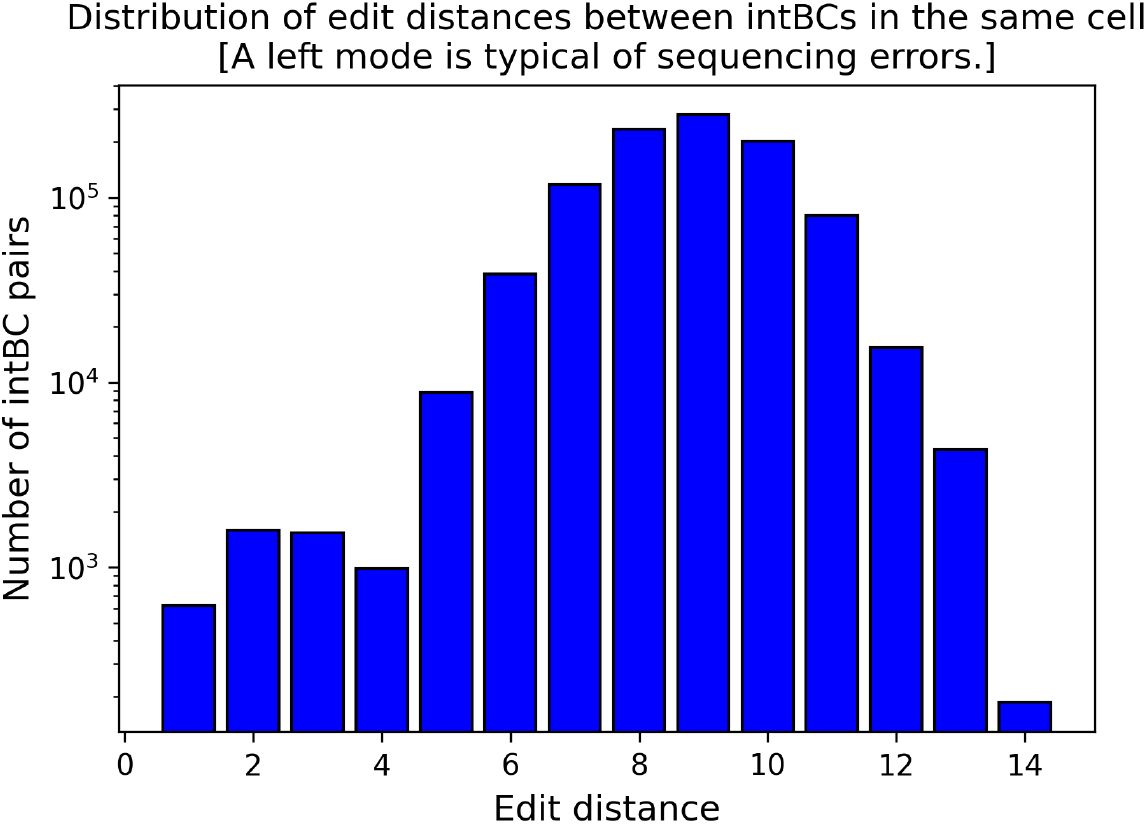
Quality control pipeline output: Histogram of edit distance distribution between intBC pairs within the same cell. The x-axis shows edit distance and the y-axis shows the number of `intBC` pairs observed in the same cell. The left mode is typical of sequencing errors; in this example the left mode is at edit distances of 2–3.

**Fig. S11.**
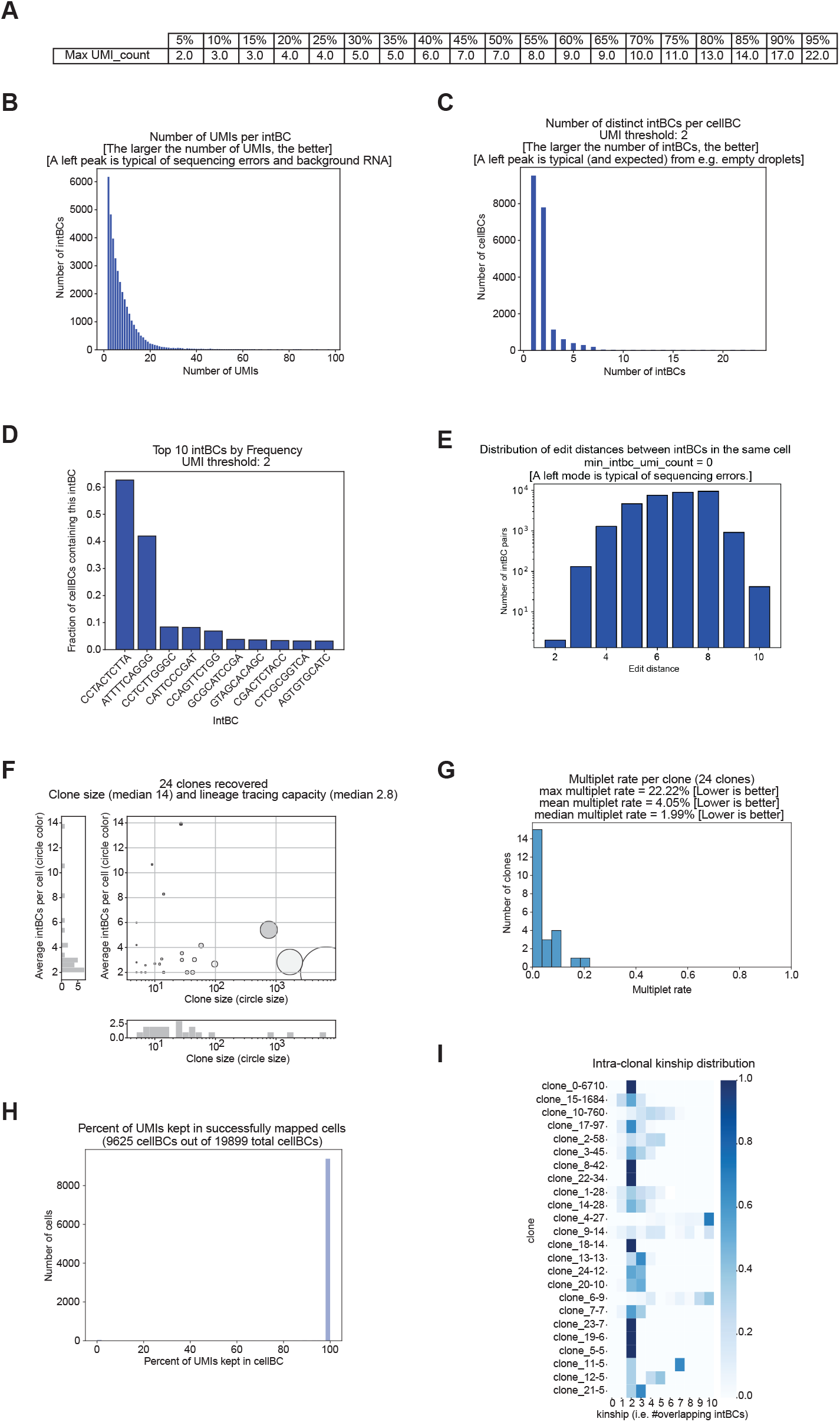
Quality control pipeline output: macsGESTALT data. (A) preQC plot-quantiles of `UMI_count`. (A) preQC plot-number of `UMI_count` per `intBC`. (C) preQC plot-number of distinct `intBCs` per `cellBC`. (D) preQC plot-top 10 frequent `intBCs` in sample. (E) preQC plot-edit distance distribution between `intBCs`. (F) postQC plot-clones sizes and average `intBCs` per `cellBC`. (G) postQC plot-multiplet rate per clone. (H) postQC plot-percentage of `UMI_count` kept in cells. (I) postQC plot-Intra-clonal kinship.

**Fig. S12.**
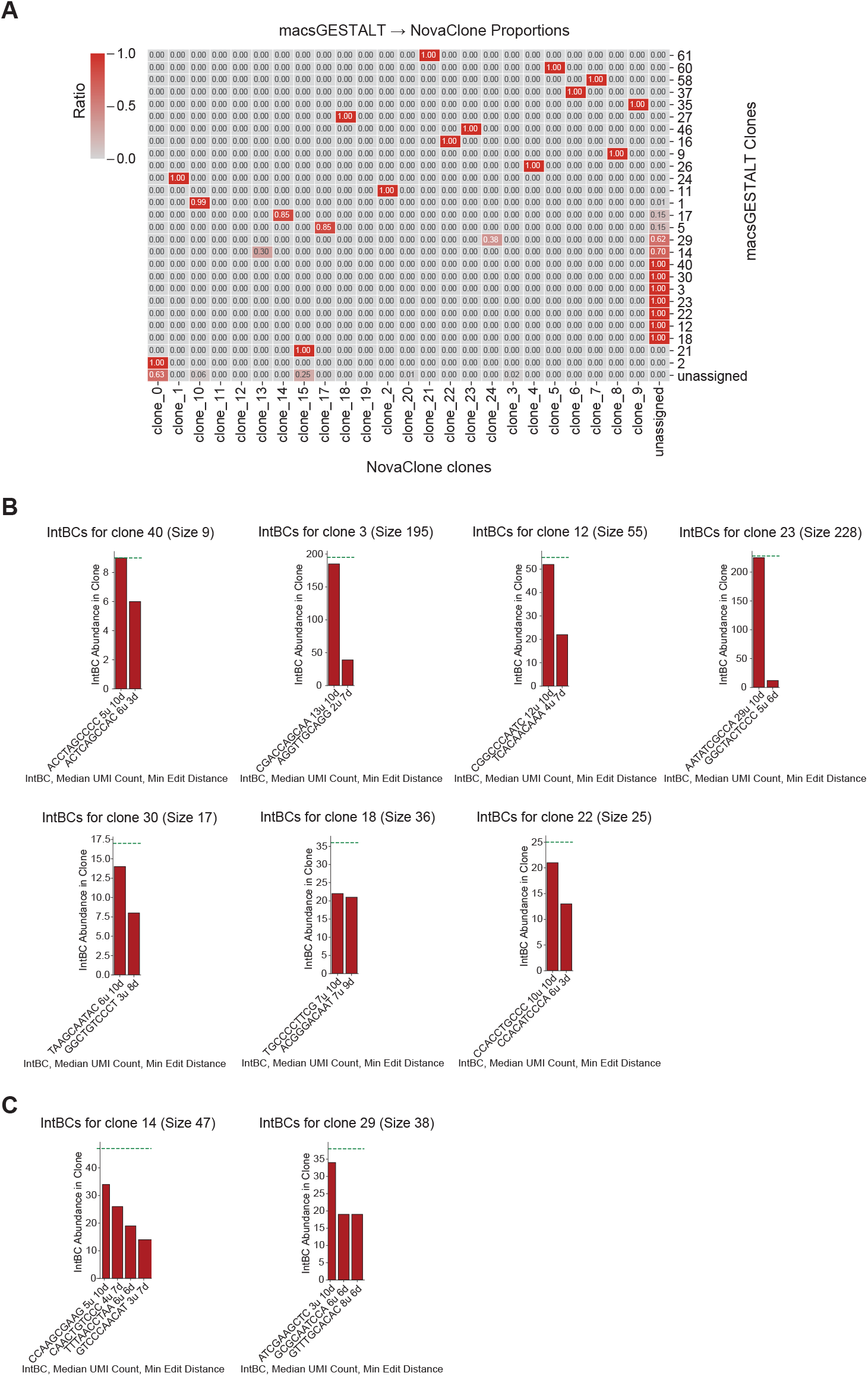
macsGESTALT clones not recovered by NovaClone. (A) Heatmap showing the proportion of `cellBCs` from each macsGESTALT clone found in `NovaClone` clone calls. Colors and annotations represent the fraction of overlap for each pair. (B) Bar plots showing the abundance of `intBCs` in macsGESTALT clones that were not recovered by `NovaClone`. The green line indicates the total number of `cellBCs` in each clone. X-axis labels display the `intBC` sequence, mean `UMI_count` and minimum edit distance from the other nearest `intBC`. (C) Bar plots showing the abundance of `intBCs` in macsGESTALT clones that were only partially recovered in `NovaClone` analysis. The green line indicates the total number of `cellBCs` in each clone. X-axis labels display the `intBC` sequence, mean `UMI_count` and minimum edit distance from the other nearest `intBC`.

**Fig. S13.**
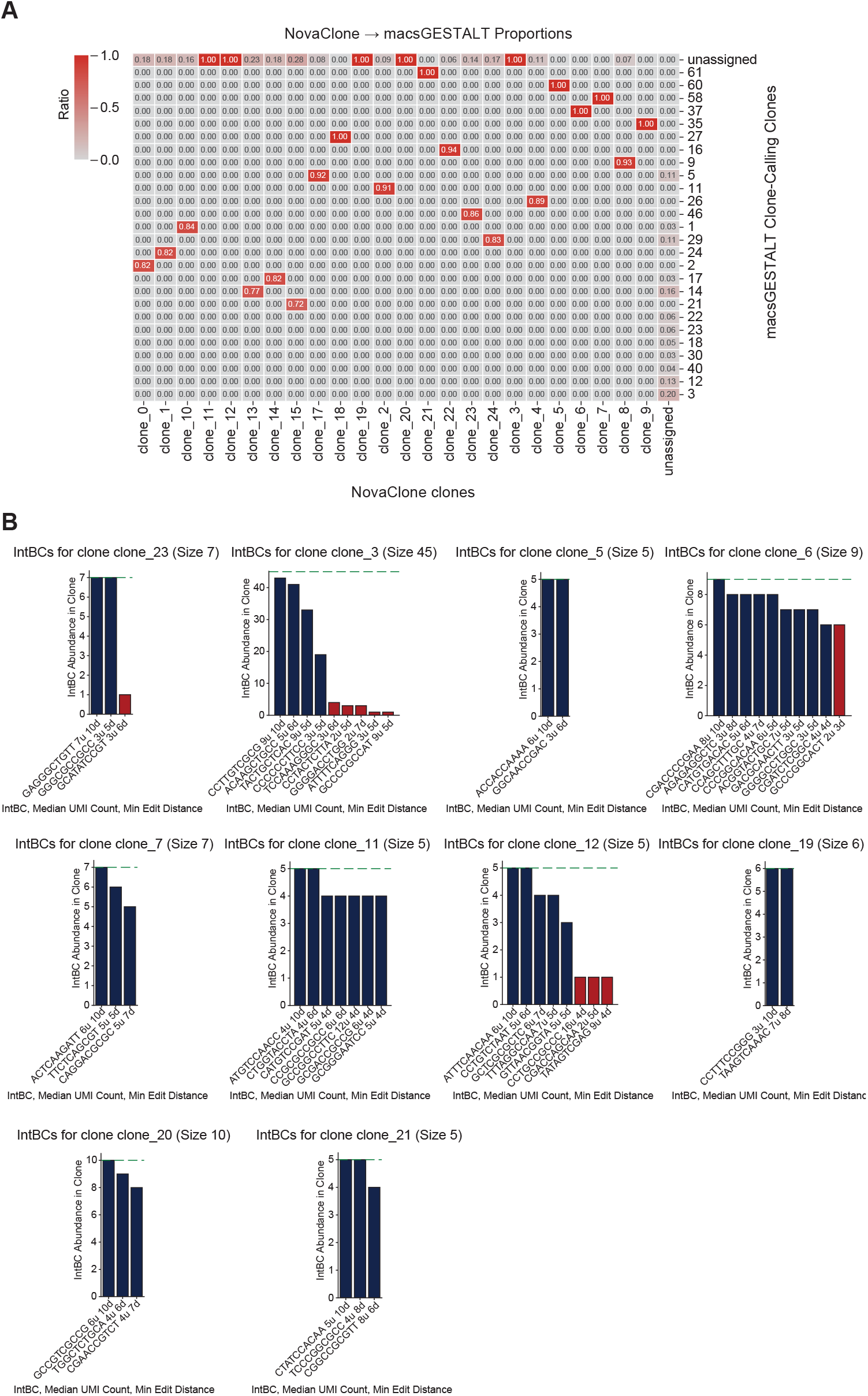
Clones recovered by NovaClone but not by macsGESTALT. (A) Heatmap showing the proportion of `cellBCs` from `NovaClone` clones found in each macsGESTALT clone. Colors and annotations represent the overlap fraction. (B) Bar plots showing the abundance of `intBCs` in clones uniquely recovered by `NovaClone`. The green line indicates the total number of `cellBCs` in each clone. X-axis labels display the `intBC` sequence, mean `UMI_count` and minimum edit distance from the other nearest `intBC`.

